# Reward-based invigoration of sequential reaching

**DOI:** 10.1101/2020.06.15.152876

**Authors:** Sebastian Sporn, Xiuli Chen, Joseph M Galea

**Affiliations:** School of Psychology and Centre for Human Brain Health, University of Birmingham, B15 2TT, UK

## Abstract

Seeking reward is a powerful tool for shaping human behaviour. While it has been demonstrated that reward invigorates performance of simple movements, its effect on more complex sequential actions is less clear. In addition, it is unknown why reward-based improvements for discrete actions are transient, i.e. performance gains are lost once reward is removed, but appear long lasting for sequential actions. We show across three experiments that reward invigorates sequential reaching performance. Driven by a reward-based increase in speed, movements also exhibited greater coarticulation, smoothness and a closer alignment to a minimum jerk trajectory. Critically, these performance gains were maintained across multiple days even after the removal of reward. We propose that coarticulation, the blending together of sub-movements into a single continuous action, provides a mechanism by which reward can invigorate sequential performance whilst also increasing efficiency. This change in efficiency appears essential for the retention of reward-based improvements in motor behaviour.

## Introduction

From drinking a cup of coffee to cleaning your teeth, sequential actions form a fundamental component of our daily life activities. When first encountered, these actions are often executed as a series of distinct sub-movements with pronounced stop periods between them ^1–8^. However, with learning these discrete sub-movements are gradually blended together to form a continuous action that is executed with increased speed, smoothness and energetic-efficiency ^9–12^. This process, known as coarticulation, is an essential mechanism for understanding skilled sequential performance acquisition as it reflects the evolution of behaviour, both temporally and spatially, towards increased efficiency ^7–12^. Coarticulation is observed across many motor behaviours such as speech production ^13,14^, sign language ^15^, piano playing ^16^, typing ^17–19^ and various other upper limb actions ^20–25^. Within sequential reaching, the coarticulation of sub-movements leads to the gradual development of a new motor primitive that is globally planned and, once initiated, must run to completion ^10^. Although this process can take weeks of practise, these newly formed motor primitives are highly generalizable and are not simply the result of increased movement speed ^10^.

It is important to emphasize that coarticulation represents a fundamentally different process from chunking which appears to feature similar characteristics. Chunking refers to the production of a series of discrete movements (i.e. button presses) that are temporally aligned ^21,22,26–28^. This is represented at a behavioural level through shorter reaction times between actions ^7,29^, however each action is still performed discretely with a pronounced stop period between them ^21,30^. In contrast, coarticulation reflects the merging of neighbouring movements into a continuous and kinematically distinct motor primitive ^9–12^.

Despite coarticulation being important to the development of skilled sequential behaviour, our ability to facilitate this process is limited ^9–11,31^. In addition, the continuous actions that arise from coarticulation appear to breakdown in several motor disorders ^32,33^. For instance, stroke patients suffering motor impairments produce actions that are decomposed into jerky sub-movements, with successful recovery being associated with a return to smooth continuous actions ^34–37^. Therefore, identifying a mechanism which can facilitate coarticulation could have far reaching implications across health and disease.

Seeking reward is a powerful tool for shaping behaviour ^38,39^. For example, the expectation of reward causes individuals to perform saccadic and reaching actions with greater speed and accuracy ^40–47^. As a result, there has been significant interest in the potential of using reward to enhance motor behaviour in both healthy individuals and clinical populations such as stroke patients ^48,49^. However, it has been shown that these reward-based effects on movement are cognitively demanding, energetically-inefficient and transient i.e. the effects are lost when reward is removed ^40–42^. Yet, these results stem from simple tasks that involve singular discrete actions e.g. movement towards a single static target. In contrast, during more complex sequential or continuous tasks, the beneficial effects of reward appear long lasting and persist even after the removal of reward ^50–52^. The mechanism, however, by which reward induces these long-lasting effects and the reason why sequential behaviour seems critical is unknown. An interesting possibility is that reward-based invigoration is associated with increased coarticulation, a mechanism which enables sequential movements to be performed faster but with greater efficiency and therefore promotes long-term retention of these performance benefits.

## Results

Here we tested this hypothesis, namely that reward can facilitate coarticulation and thereby promote energetically efficient performance gains that persist even in the absence of reward. To this end, our main experiment (N=42) assessed the effect of reward on sequential reaching performance and the evolution of coarticulation over the course of two testing days. We then carried out two further experiments to assess the robustness of these performance gains during an additional testing day without reward availability (N=5), and to investigate whether reward or performance-based feedback drove these observed improvements (N=60).

We developed a novel reaching task in which participants made 8 sequential reaching movements to designated targets (1 trial) using a motion tracking device (Figure 1a,b). Participant behaviour could range from executing 8 individual movements (i.e., stopping at each via point; Figure 1c) to a series of 5 coarticulated movements that would reflect the outcome of minimising jerk across the trial (Methods Equation 3; Figure 1d) ^12^. Changes in the number of reaching movements were associated with marked differences in the velocity profile. Specifically, individual reaches were characterised by pronounced stops in-between movements, with velocity dropping close to zero. As reaching movements merge through coarticulation, these dips in velocity gradually disappear (Figure 1c). As suggested by the minimum jerk model (Methods, Equation 3), we hypothesised that coarticulation would mainly occur in the central set of three in-centre-out reaching movements (Figure 1c,d) ^9–12,53^.

**Figure 1.**
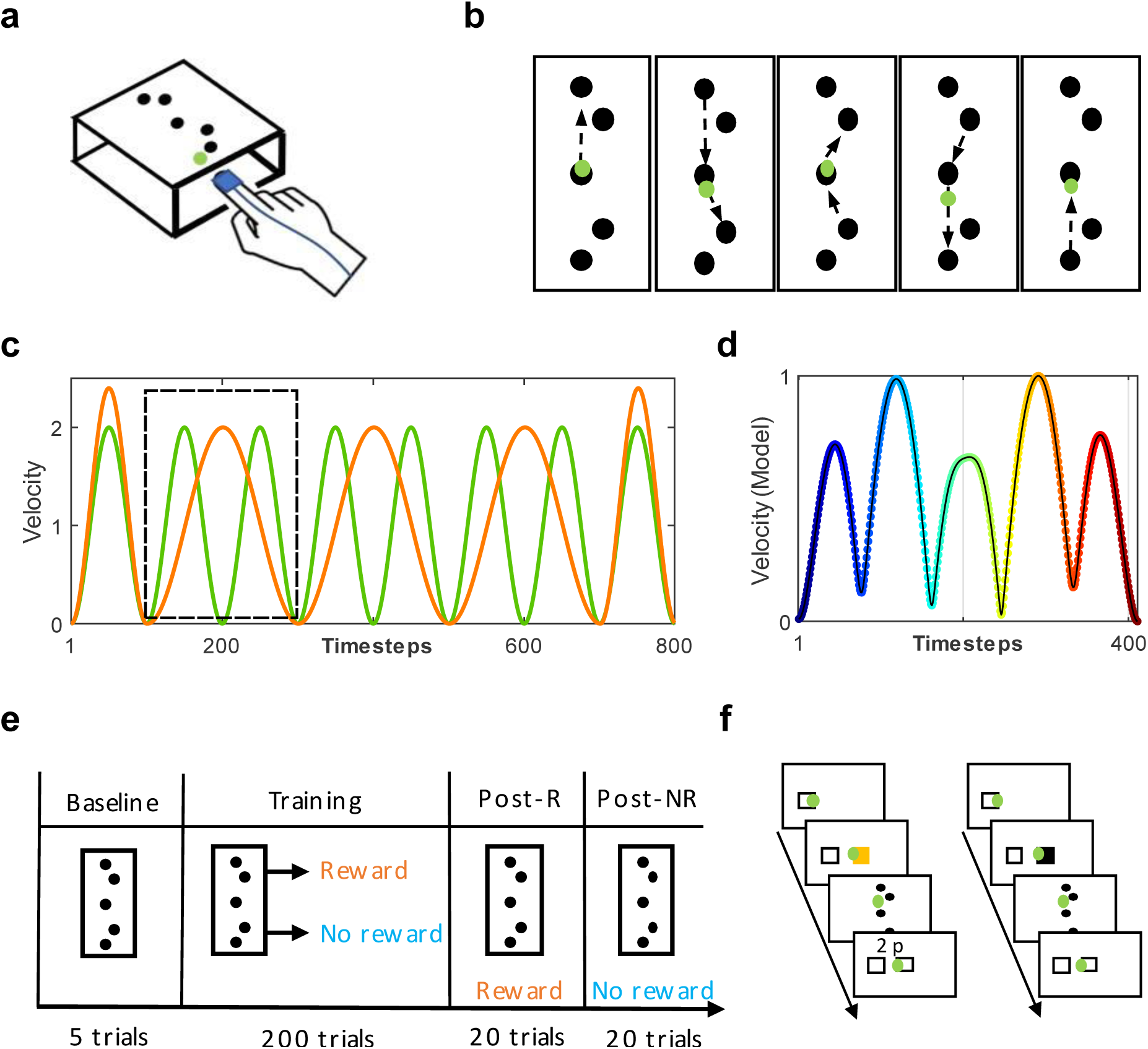
Experimental setup. **a)** Participants wore a motion-tracking device on the index finger and the unseen reaching movements were performed across a table whilst a green cursor matching the position of index finger was viewable on a screen. **b)** 8 movement sequential reaching task. The participants started from the centre target. **c)** Schematic velocity profiles highlighting the difference between single versus coarticulated reaching movements for a trial. Based on the predictions of a minimum jerk model, we assumed that the three sets of in-centre-out reaching movements could be coarticulated which would reduce the number of velocity peaks from 8 (green) to 5 (orange). **d)** Velocity profile for a trial predicted by a minimum jerk model. **e)** Study design. Prior to the start of the experiment, participants were trained on the reaching sequence and were then asked to perform 10 baseline trials. Randomly allocated to a reward and no reward group, participants completed 200 trials during training and an additional 20 trials in each post assessment; one with reward (post-R) and one without (post-NR) (counterbalanced across participants). This design was repeated 24 hours later (Day 2). **f)** Rewarded trials were cued using a visual stimulus prior to the start of the trial. At the end of the trial, participants received trial-based monetary feedback based on their last 20 trials (close-loop design). In no reward trials, participants were instructed to be as fast and accurate as possible.

Participants were randomly allocated to a reward or no reward group. Prior to the start of the experiment, participants were trained on the sequence without a time constraint until reaching a learning criterion of 5 successful trials in a row. Therefore, any performance gains could only be attributed to improvements in the execution and not memory of the sequence. There was then a baseline period (10 trials), during which both groups were encouraged to complete each trial “as fast and as accurately as possible”. Afterwards, during training, both groups completed 200 training trials. Participants in the reward group were able to earn money depending on their movement time (MT) and received performance-based feedback (the amount of money awarded in a given trial) after reaching the last target in each trial. Reward trials were cued with a yellow start box (Figure 1f) and the monetary reward was calculated using a closed-loop design comparing MT performance on a given trial with performance on the last 20 trials. Participants in the no reward group were instructed to move as fast and accurately as possible. Importantly, participants did not receive feedback on their level of coarticulation. Missing a target resulted in an immediate abortion of the current trial, which they then had to repeat, thereby forcing participants to accurately hit each target. After training, both groups engaged in a rewarded (post-R) and no rewarded (post-NR) post-assessment (20 trials each). The post-assessment intended to compare performance between groups when under the same condition (Figure 1e). This design was repeated 24 hours later.

### Reward invigorates sequential reaching movements

Movement time (MT) reflected the total movement duration from exiting the start box until reaching the last target. Our results highlight that reward instantaneously invigorates sequential reaching behaviour with these performance gains being maintained even in the absence of reward (Figure 2a). Specifically, we found a significant decrease in MT over the course of training on both days for both groups (mixed-effect ANOVA; timepoint (early (1^st^ 20 trials) vs late (last 20 trials)) x group; main effect for timepoint, day 1: F = 18.29, p < 0.0001; day2: F = 8.51, p = 0.0058; Figure 2b,d). Importantly, despite no differences in MT between groups during baseline on day 1 (Wilcoxon test; Z = −1.38, p = 0.17), we found that the reward group produced significantly faster MTs than the no reward group across training on both days (main effect for group, day 1: F = 18.96, p < 0.0001; day 2: F = 17.58, p < 0.0001; Figure 2b,d).

**Figure 2.**
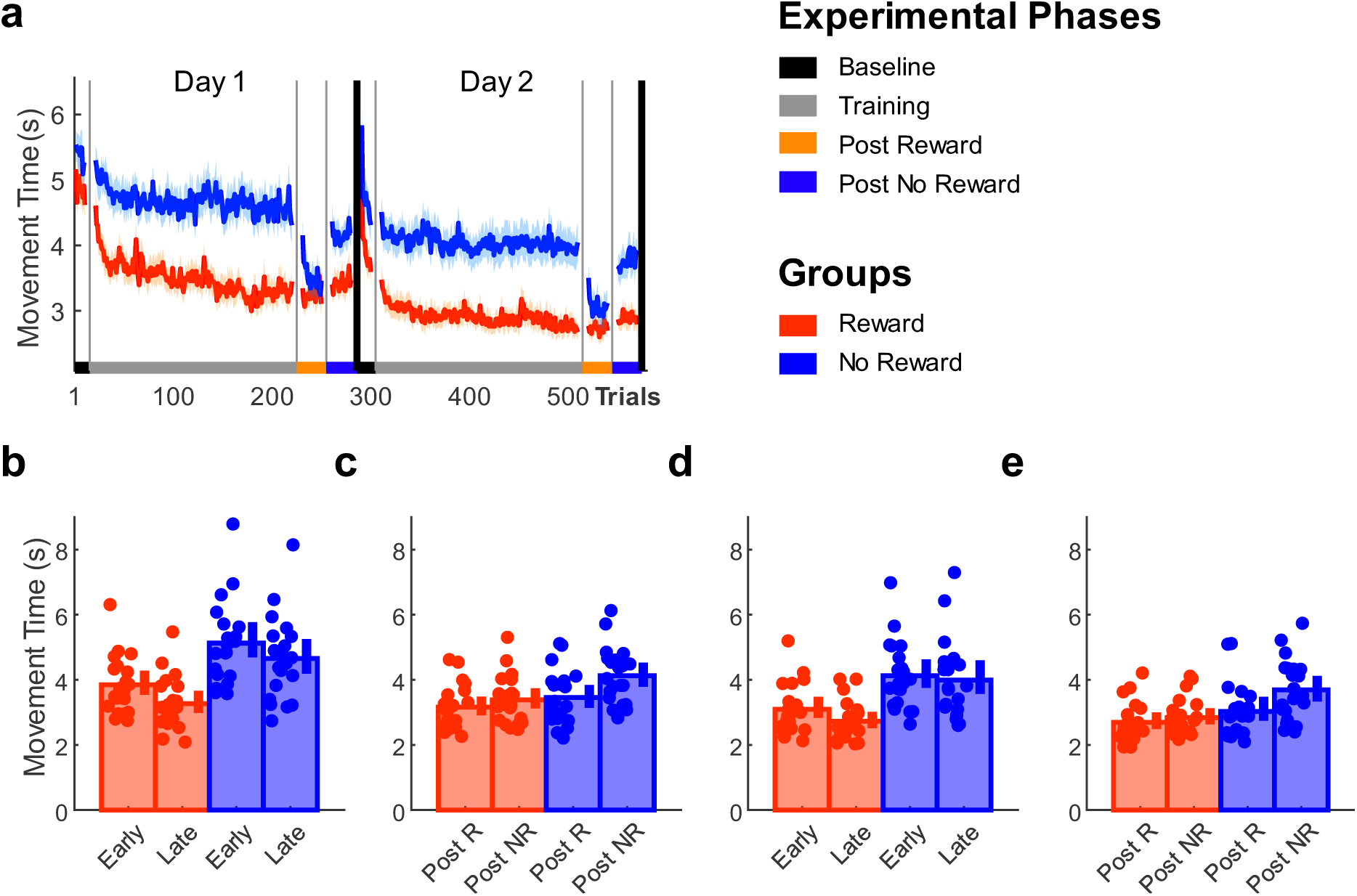
Reward-based improvements in MT. **a)** Trial-by-trial changes in MT averaged over participants for both groups. **b-e)** Median MT values for each participant for both groups. **b)** Comparing MT performance during training on day 1 (early (first 20 trials) vs late (last 20 trials)). **c)** Post assessment performance (day 1; post-R vs post-NR). **d)** Training (day 2; early vs late). **e)** Post assessment performance (day 2; post-R vs post-NR)

Across post assessments, a significant interaction was found between phase (post-R (all 20 trials) vs post-NR (all 20 trials)) and group (mixed ANOVA; condition x group; interaction, day 1: F = 18.07, p < 0.0001; day 2: F = 19.99, p < 0.0001). Specifically, there was a significant difference in MT between groups during post-NR (Wilcoxon test; day 1: Z = −2.82, p = 0.0192; day 2: Z = −3.27, p = 0.0044; Figure 2c,e) but not post-R (Wilcoxon test; day 1: Z = −1.13, p = 1; day 2: Z = −1.38, p = 1). This indicates that the no reward group were able to instantaneously invigorate their performance during post-R. However, these performance gains were not maintained during post-NR, suggesting that they remained transient in nature. In contrast, the enhanced MT performance in the reward group did not change significantly between post assessments, implying that performance gains had become reward-independent (Wilcoxon test; day 1: Z = −1.11, p = 1; day 2: Z = −0.91, p = 1; Figure 2c,e). The counterbalancing of post assessment order did not affect these results (Supplementary Figure 1).

These decreases in MT could be driven by two processes: (1) an invigoration of each reaching movements’s peak velocity which has been observed to underlie the transient performance gains in singular discrete reaching tasks ^40–42^ and (2) the coarticulation of sequential reaching movements which decreases MTs via a reduction in dwell time around the central target.

### Reward-based invigoration of peak velocities is instantaneous

To assess changes in peak velocities (Methods, equation 1), we averaged peak velocity across the 8 reaching movements (Figure 3a). Over the course of training, we found significant increases in peak velocity on both days (mixed ANOVA; timepoint x group; main effect for timepoint, day 1: F = 9.72, p = 0.0034; day 2: F = 8.56, p = 0.0057; Figure 3b,d). Despite there being no differences in peak velocities during baseline on day 1 (Wilcoxon test; Z = 0.70, p = 0.4812), the reward group produced significantly higher peak velocities than the no reward group across training (main effect for group, day 1: F = 19.42, p < 0.0001; day 2: F = 19.28, p < 0.0001; Figure 3b,d). This supports existing findings that reward-based invigoration of MT can be driven by increases in peak velocities ^40–42^. Yet again, the no reward group exhibited a pronounced “on-off” effect across post assessments (post-R vs post-NR) for both days (mixed-effect ANOVA; condition x group; interaction, day 1: F = 8.02, p = 0.0072; day 2: F = 24.92, p < 0.0001). Specifically, we found significant differences between groups during post-NR (Wilcoxon test; day 1: Z = 2.84, p = 0.018; day 2: Z = 3.07, p = 0.0084; Figure 3c,e) but not post-R (Wilcoxon test; day 1: Z = 0.86, p = 1; day 2: Z = 0.93, p = 1; Figure 3c,e). Similarly to MT, these results suggest that the no reward group were able to instantaneously increase peak velocity during post-R, but these performance gains remained transient in nature and were not maintained during post-NR. In contrast, peak velocities in the reward group remained elevated across post assessments irrespective of reward availability (Wilcoxon test; day 1: Z = 0.86, p = 1; day 2: Z = 0.68, p = 1; Figure 3c,e).

**Figure 3.**
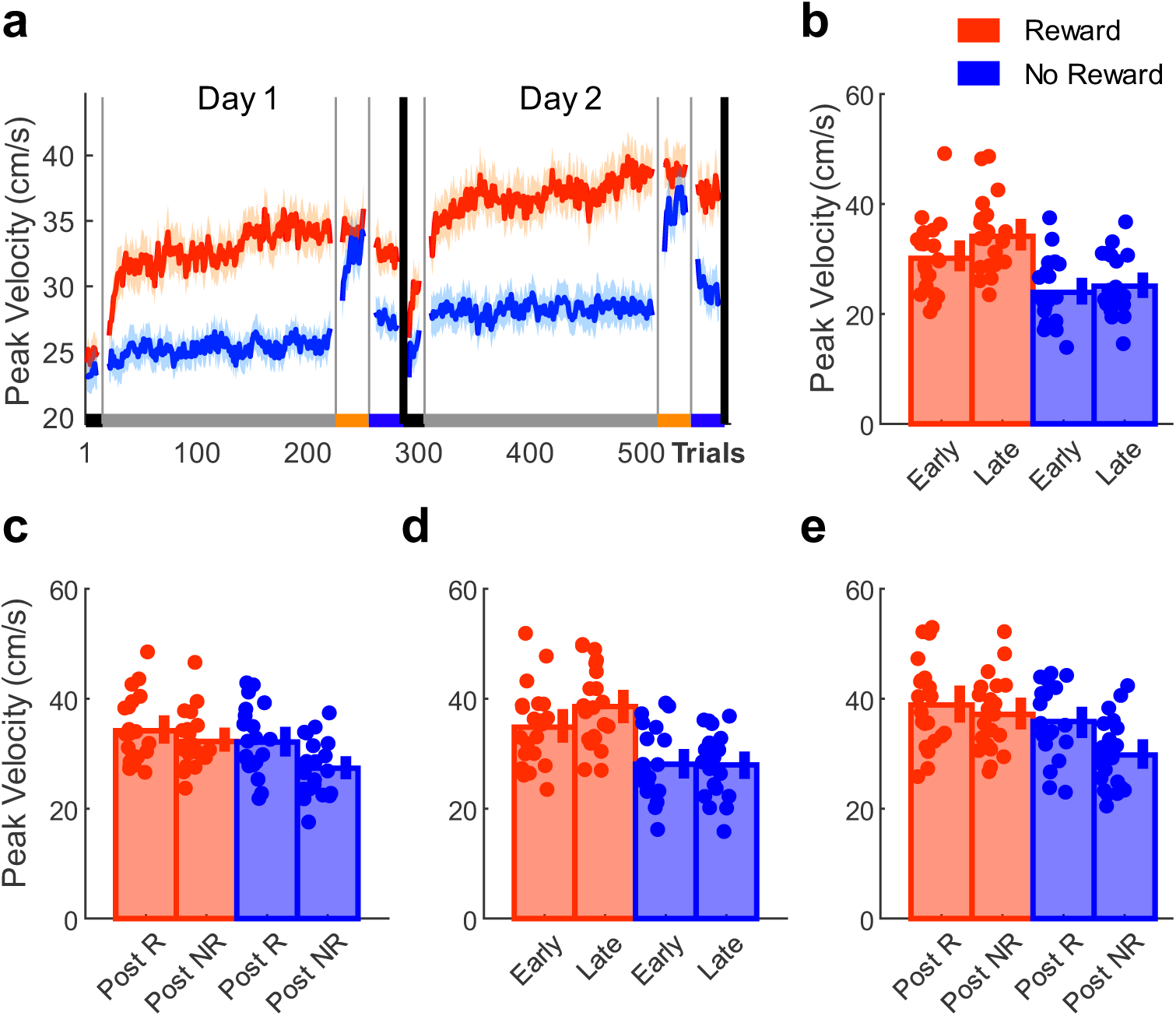
Reward-based improvements in peak velocity. **a)** Trial-by-trial changes in peak velocity averaged over participants for both groups. **b-e)** Median peak velocity values for each participant for both groups. **b)** Comparing peak velocity performance during training on day 1 (early (first 20 trials) vs late (last 20 trials)). **c)** Post assessment performance (day 1; post-R vs post-NR). **d)** Training (day 2; early vs late). **e)** Post assessment performance (day 2; post-R vs post-NR)

### Reward-based facilitation of coarticulation is training-dependent

Coarticulation describes the blending of individual motor elements into a combined smooth action. This is represented in the velocity profile by the stop period between two movements gradually disappearing and being replaced by a single velocity peak (Figure 1c). To measure coarticulation, we developed a coarticulation index (CI; Methods, Equation 2) that compared the mean peak velocities of two sequential reaches with the minimum velocity around the via point. The smaller the difference between these values, the greater coarticulation had occurred of these two movements as reflected by CI value closer to 1 (Figure 4a). As the central 3 segments of the movement could potentially be coarticulated the CI ranged from 0-3 for each trial (Figure 4b). Our results show that reward facilitates movement coarticulation and leads to stable changes in behaviour even in the absence of reward. Additionally changes in coarticulation, in contrast to the changes in peak velocities, appears to be training-dependent. Across training, CI levels increased on both days (mixed ANOVA; timepoint x group; main effect for timepoint, day 1: F = 21.70, p < 0.0001; day 2: F = 21.45, p < 0.0001; Figure 4b). Crucially, although there were no differences between groups in CI levels during baseline (Wilcoxon test; Z = 1.31, p = 0.1908), we found significantly higher CI levels for the reward group on both days (main effect for group, day 1: F = 6.81, p = 0.0127; day 2: F = 9.10, p = 0.0044; Figure 4c,e). However, unlike MT and peak velocity, no significant increases in CI levels were observed for the no reward group on either day during post-R. This suggests that coarticulation cannot be invigorated instantaneously but represents a training-dependent process that is facilitated by reward. Importantly, CI levels were maintained during post-NR on both days for the reward group, highlighting that changes in coarticulation had become reward-independent (mixed-effect ANOVA; condition x group; main effect for group, day 1: F = 4.91, p = 0.0324; day 2: F = 7.38, p = 0.0097; Figure 4d,f). To understand whether CI levels are related to the retention of MT performance, we correlated MT values with CI levels during post-NR (Figure 4g,h). We found a significant correlation between CI levels and MT performance for both day 1 (partial correlation controlling for group; post-NR: ρ = −0.51; p < 0.0001, Figure 4g) and day 2 (partial correlation controlling for group; post-: ρ = - 0.66, p < 0.0001, Figure 4h). Although not causal, this indicates that faster MTs during post-NR were associated with higher levels of coarticulation and that the reward group showed better performance for both (Figure 4g,h).

**Figure 4.**
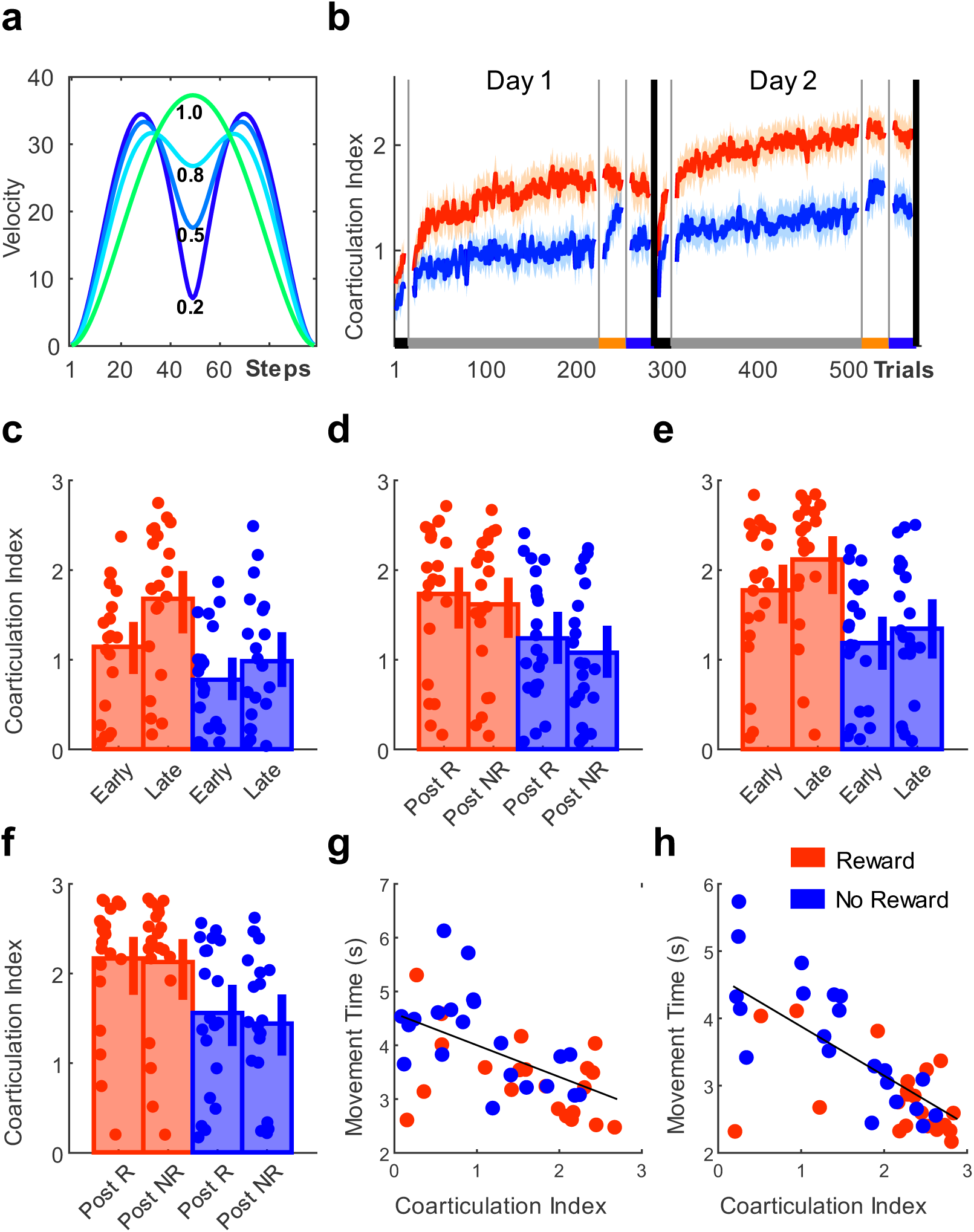
Reward-based improvements in CI levels. **a)** Illustration of CI levels. **b)** Trial-by-trial changes in CI levels averaged over participants for both groups. **c-f)** Median CI levels for each participant for both groups. **c)** Comparing CI levels during training on day 1 (early (first 20 trials) vs late (last 20 trials)). **d)** Post assessment performance (day 1; post-R vs post-NR). **e)** Training (day 2; early vs late). **f)** Post assessment performance (day 2; post-R vs post-NR). **g-h)** Scatterplots displaying the relationship between MT and CI levels during post-NR on **g)** day 1 and **h)** day 2 with a linear line fitted across groups.

To summarise, these results demonstrate that improvements in MT are driven by two processes which are both reward-sensitive but follow different time courses. The invigoration of peak velocities is instantaneous, whereas coarticulation is training-dependent. Invigoration in the no reward group was mainly driven by increases in peak velocities which were transient In nature, thereby displaying a pronounced “on-off” effect. In contrast, participants in the reward group capitalised on both strategies: the invigoration of peak velocities and additionally increases in coarticulation which led to persistent and ultimately reward-independent improvements in MT performance. Importantly, despite an increase in execution errors on day 1 (reward group), we found that execution errors were not associated with performance levels during late training (day 2; Supplementary Figure 2). In addition, no significant differences between groups were found on day 2 (mixed-effect ANOVA; condition x group; main effect for group, day 2: F = 1.88, p = 0.1785), suggesting that performance gains by the second testing day reflected true improvements in skill.

### Reward-based improvements in smoothness are associated with reward-independent maintenance of performance gains

It has been suggested that coarticulation reduces motor costs via the reduction of jerk and thereby reflects more efficient (smoother) execution ^53,54^, which in turn could explain why participants in the reward group maintained their improved performance levels even in the absence of reward. Using spectral arc length as a smoothness metric^55^, we show that the performance in the reward group became progressively smoother which could reflect increased energetic efficiency (Figure 5a) ^54^.

**Figure 5.**
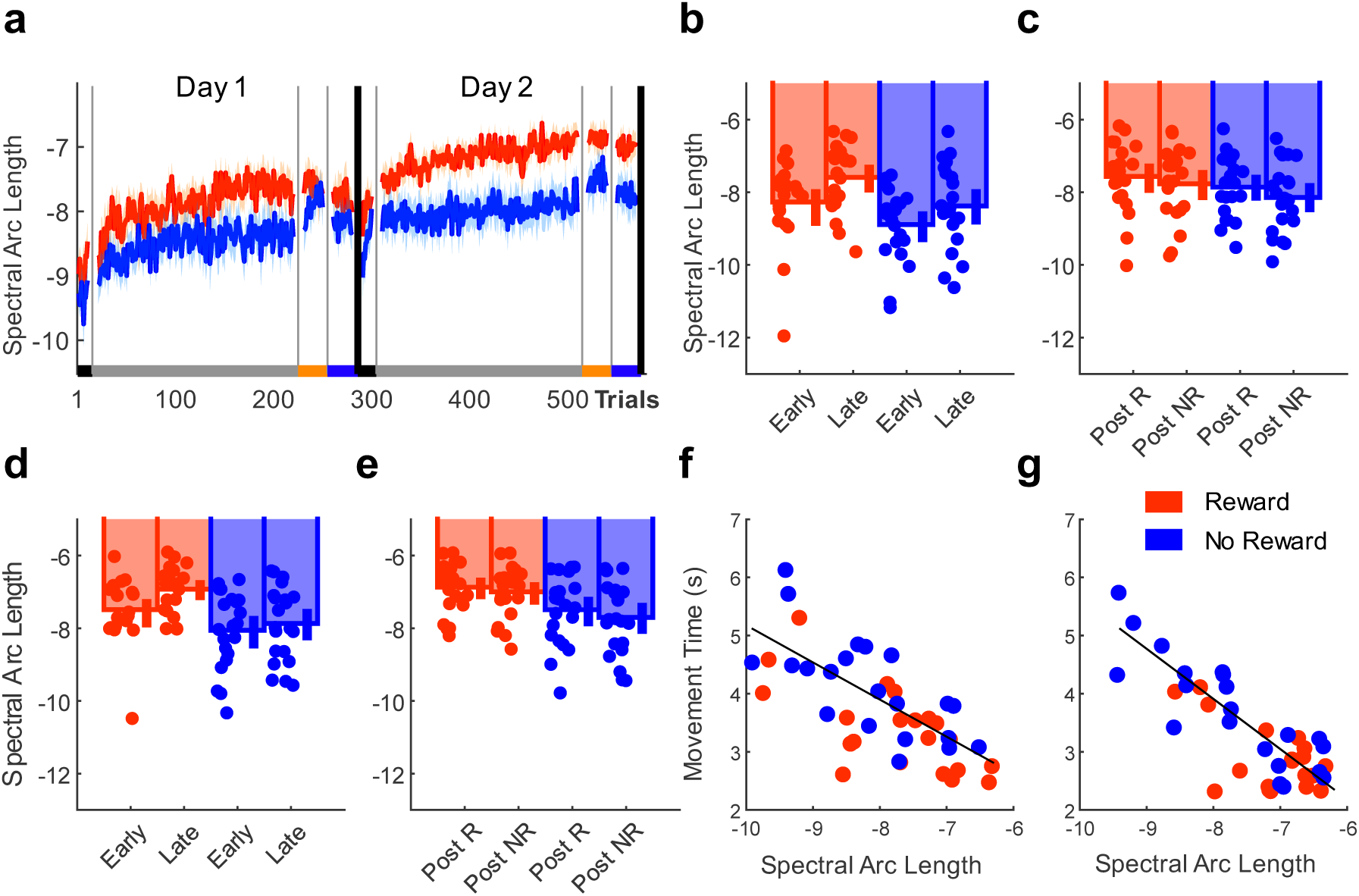
Reward-based improvements in smoothness. **a)** Trial-by-trial changes in spectral arc length (smoothness) averaged over participants for both groups. A value closer to zero indicates greater smoothness^55^. **b-e)** Median smoothness values for each participant for both groups. **b)** Comparing smoothness during training on day 1 (early (first 20 trials) vs late (last 20 trials)). **c)** Post assessment performance (day 1; post-R vs post-NR). **d)** Training (day 2; early vs late). **e)** Post assessment performance (day 2; post-R vs post-NR). **f-g)** Scatterplots displaying the relationship between MT and smoothness during post-NR on **f)** day 1 and **g)** day 2 with a linear line fitted across groups.

Movement smoothness significantly improved across training on both days (mixed ANOVA; timepoint x group; main effect for timepoint, day 1: F = 27.56, p < 0.0001; day 2: F = 18.19, p < 0.0001, Figure 5a). Despite there being no differences between groups during baseline (Wilcoxon test; Z = 1.79, p = 0.0741), the reward group showed a greater improvement in smoothness throughout training (main effect for group, day 1: F = 5.31, p = 0.027; day 2: F = 7.78, p = 0.0080; Figure 5b,d). During post assessments, we found a significant group effect for day 2 (mixed-effect ANOVA; post assessment x group; group, day 1: F = 1.44, p = 0.2369; day 2: F = 6.41, p = 0.0154; Figure 5c,e). This suggests that improvements in smoothness were greater in the reward group and became reward-independent. Additionally, we found that increased smoothness was strongly associated with faster MTs when reward was not available. We correlated smoothness values with MTs during post-NR on both days (Figure 5f,g). We found a significant correlation between smoothness and MT performance for both day 1 (partial correlation controlling for group; Post-NR: ρ = - 0.69; p < 0.0001, Figure 5f) and day 2 (partial correlation controlling for group; post-: ρ = - 0.79, p < 0.0001, Figure 5g). These results align with the aforementioned results showing that CI levels were associated with faster MTs during post-NR, and point to improvements in movement efficiency as a potential mechanism enabling reward-independent maintenance of performance gains during post-NR.

### Progressive alignment to the predictions of a minimum jerk model is facilitated by reward

We then assessed whether performance aligned with the predictions of a minimum jerk model ^12^ by calculating the mean squard error between the model’s and the actual velocity profile on a trial-by-trial basis (Methods, Equation 3; Figure 6a). In comparison to the no reward group, performance in the reward group became significantly more aligned to the predictions of the minimum-jerk model. Mean squared error progressively decreased across days although this was not significant for day 2 (mixed ANOVA; timepoint x group; main effect for timepoint, day 1: F = 16.19, p < 0.0001; day 2: F = 2.23, p = 0.1429, Figure 6b). Despite no differences between groups during baseline (Wilcoxon test; Z = −1.16, p = 0.2472), the reward group’s performance showed significantly greater similarity to the model’ s predictions (main effect for group day 1: F = 5.61, p = 0.0228; day 2: F = 7.56, p = 0.0089, Figure 6c,e). Across post assessments, we found a significant interaction for day 2 (mixed-effect ANOVA; condition x group; main effect for interaction, for day 1: F = 3.92 p = 0.0548; day 2: F = 7.58, p = 0.0088, Figure 6b,f), however all post-hoc comparisons were not significant. Nevertheless, there was a significant group effect for day 2 (main effect for group, for day 1: F = 3.26 p = 0.0787; day 2: F = 4.98, p = 0.0313, Figure 6b,f), suggesting that the degree of similarity to the minimum-jerk model was maintained across post assessments irrespective of reward availability.

**Figure 6.**
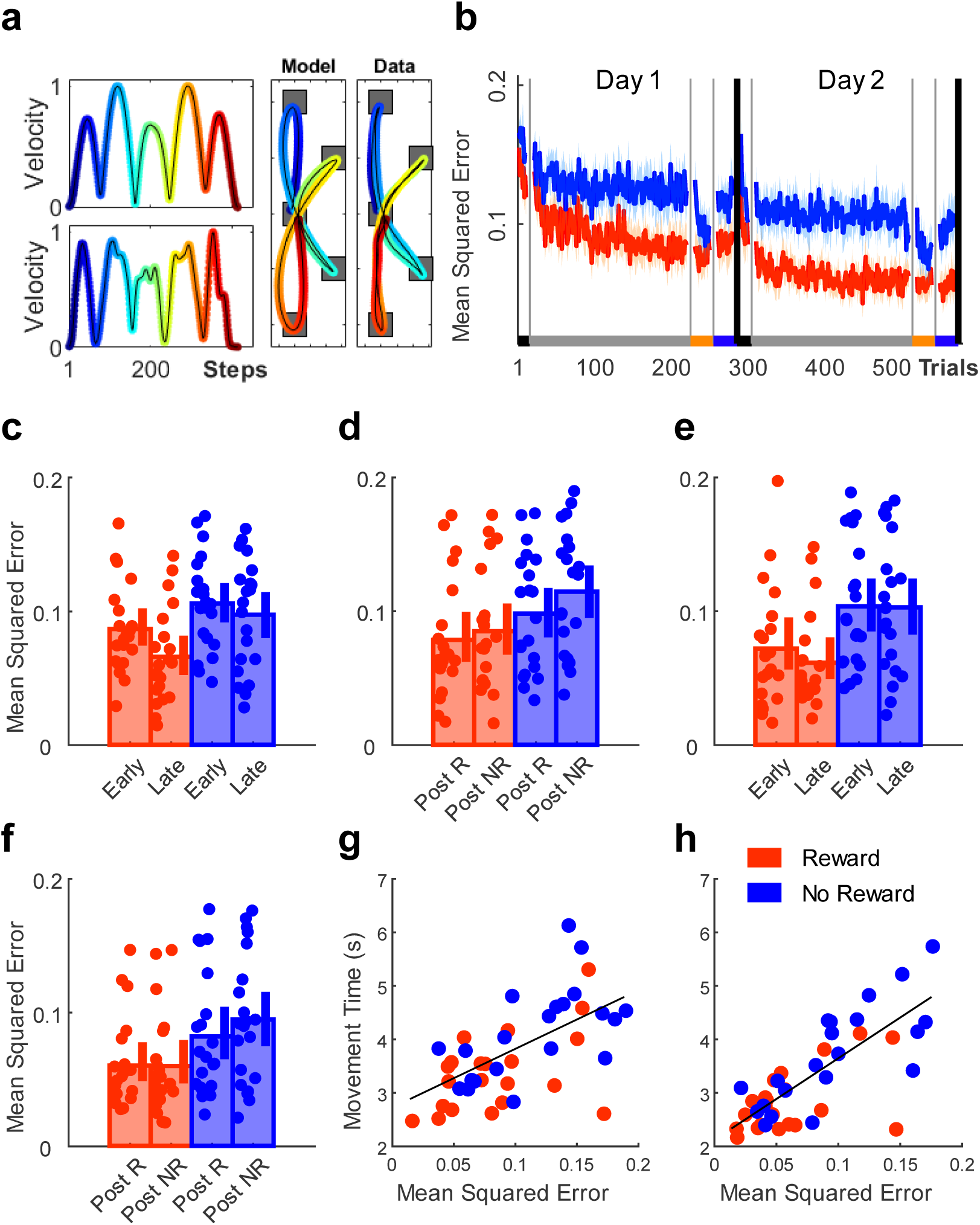
Progressive alignment to the predictions of a minimum jerk model is facilitated by reward. **a)** Comparisons between data and the predictions of a minimum jerk model for both trajectory (right panel) and velocity profiles (left panel) for a single participant. **b)** Trial-by-trial changes in mean square error averaged over participants for both groups. **c-f)** Median mean square error for each participant in both groups. **c)** Day 1 early vs late training **d)** Day 1 post assessment (post-R vs post-NR). **e)** Day 2 early vs late training **f)** Day 2 post assessment. **g-h)** Scatterplots displaying the relationship between MT and mean squared error during post NR on **g)** day 1 and **h)** day 2 with a linear line fitted across groups.

Within this context, we aimed to assess whether more optimal performance with regards to the model prediction was related to the maintenance of performance. Using partial linear correlations, we found a correlation between model similarity and MT performance during post-NR for both day 1 (partial correlation controlling for group; Post-NR: ρ = 0.55; p < 0.0001, Figure 6g) and day 2 (partial correlation controlling for group; post-: ρ = 0.70, p < 0.0001, Figure 6h). These results complement our findings on smoothness and CI. Overall, our results show that over the course of the experiment performance in the reward group became smoother and more optimal with regards to the predictions of a minimum jerk model. Potentially due to being more energetically efficient ^56^, participants in the reward group maintained MT performance gains even in the absence of reward.

### Spatial reorganisation identifies the final stages of coarticulation and is enhanced by reward

Coarticulation can also be expressed spatially as the radial distance between the peak velocity on the sub-movements and the minimum velocity around the via point (Figure 7a). This distance becomes smaller with high levels of coarticulation ^9,10^ (Figure 7a; Supplementary Figure 3), suggesting that spatial reorganisation reflects the final stages of two movements merging together. We found that the reward group expressed spatial reorganisation with significant decreases in the radial distance between peaks and the via point. No difference between groups in radial distance was observed during baseline (Wilcoxon test; Z = −1.18, p = 0.2371). However, there was significant main effects for timepoint (mixed ANOVA; timepoint x group; main effect for timepoint, day 1: F = 7.32, p = 0.0100; day 2: F = 18.14, p < 0.0001; Figure 7b) as well as group for the second testing day (main effect for group, day 1: F = 1.95, p = 0.1706; day 2: F = 7.52, p = 0.0091; Figure 7c,e). This indicates that radial distance progressively decreased across both days, with the reward group showing greater changes on day 2. Across post assessments, the no reward group showed no reward-driven changes in spatial reorganisation, whereas the reward group maintained their performance although the differences between groups was only significant on day 2 (mixed-effect ANOVA; post assessment x group; main effect for group, day 1: F = 3.08, p = 0.0868; day 2: F = 5.76, p = 0.0211; Figure 7d,f).

**Figure 7.**
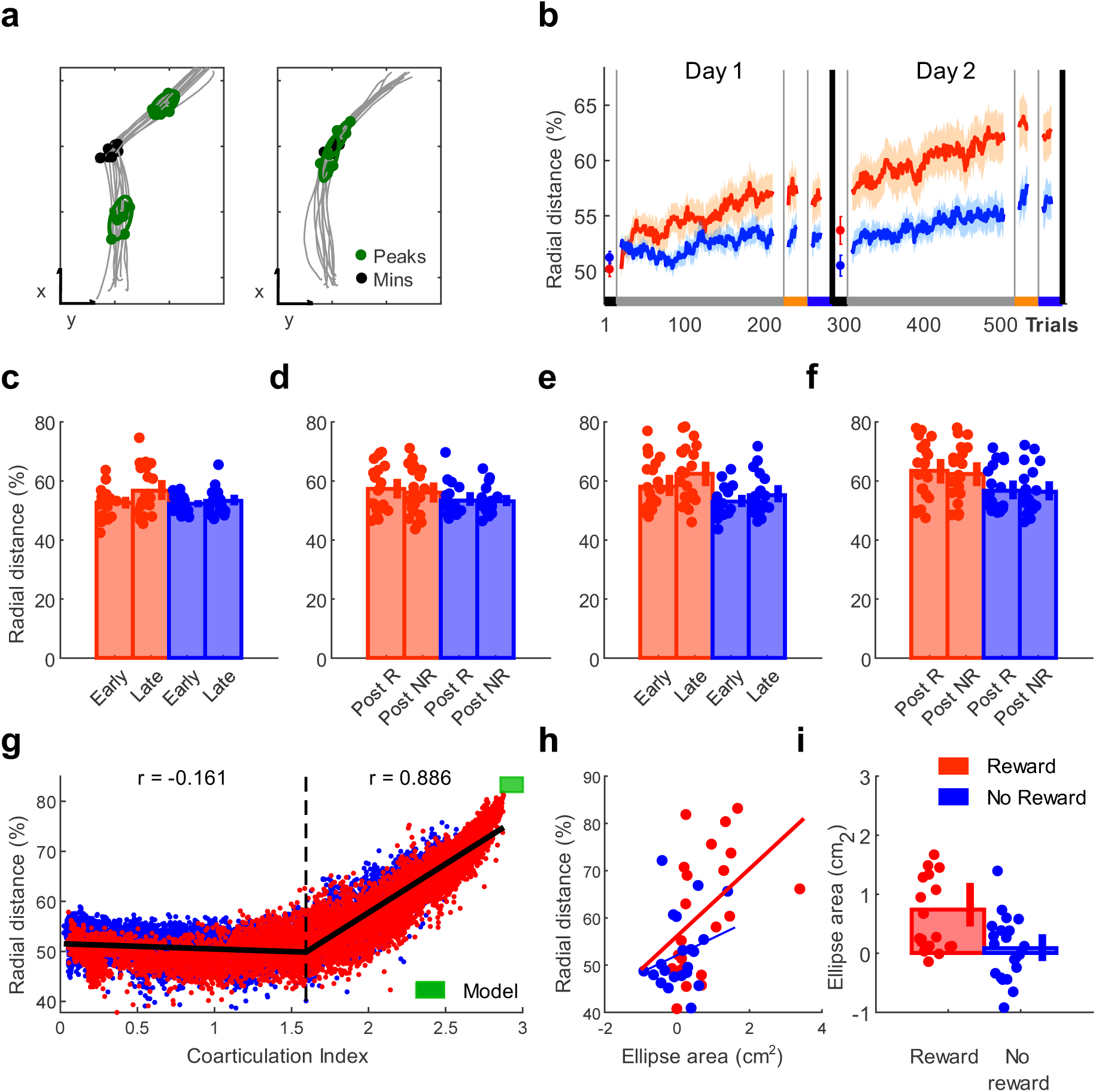
Reward-based improvements in spatial reorganisation. **a)** Example data for the spatial location of peak velocities when performing two individual (left panel) and one coarticulated (right panel) reaching movement. **b)** Trial-by-trial changes in radial distance averaged over participants for both groups. **c-f)** Median radial distance for each participant for both groups. **c)** Comparing radial distance during training on day 1 (early (first 20 trials) vs late (last 20 trials)). **d)** Post assessment performance (day 1; post-R vs post-NR). **e)** Training (day 2; early vs late). **f)** Post assessment performance (day 2; post-R vs post-NR). **g)** Scatterplot illustrating the relationship between mean CI levels and spatial reorganisation (radial distance %). It includes a two-segment piecewise linear function fitted to the data, while the green area represents the model prediction. **h)** Scatter plot displaying the relationship between variability (ellipse area cm^2^) during early training and spatial coarticulation (radial distance %) during late training. **i)** Bar plot comparing the reward and no reward group early training variability including jittered scatter of participant-based median (ellipse area cm^2^).

To understand the relationship between CI and spatial reorganisation, we plotted them against each other and detected a pronounced drift in radial distance values (%) with increasing CI levels resulting in a curvilinear shape (Figure 7g). After fitting a two-segment piecewise linear function to the data, we found an inflection point at ∼1.66 (CI) and a strong correlation between CI levels and radial distance values for the second segment (partial correlation controlling for group; segment1: ρ = −0.16; p <0.0001; segment 2: ρ = .89; p< 0.0001).

This suggests that in order to fully coarticulate two consecutive movements, spatial reorganisation is essential ^9–11^ with this process appearing significantly more pronounced in the reward group. Overall, these findings indicate that improvements in movement efficiency through a quantitative change in how the task is performed may enable the retention of reward-based performance gains.

Within the context of reinforcement learning, it has been suggested that reward leads to improvements in performance via increases in exploration during the early stages of training ^57,58^. To assess whether an early increase in spatial variability was associated with improved performance, we measured the area of the confidence ellipses used to determine the radial distance to the via point. We then correlated the ellipse area (cm^2^), which was normalised to baseline, over the first 30 trials on day 1 with radial distance (%) over the last 30 trials on day 2 (Figure 7h). We found a significant correlation (partial correlation controlling for group; ρ = .40; p = 0.0104) suggesting that early increases in spatial variability were associated with increases in spatial coarticulation towards the end of the experiment. Comparing ellipse area between groups during early training (first 30 trials), we found that the reward group exhibited higher levels of variability (Wilcoxon test, Z = 2.52, p = 0.0119, Figure 7i). These results indicate that early reward-driven increases in spatial variability may benefit future performance.

Performance gains are maintained across an additional testing day without reward

We next aimed to assess the robustness of these performance gains in an experiment including an additional testing day without reward availability (elongated washout condition). In experiment two, participants underwent the same regime as the reward group in experiment 1 on the first two days. On the third day, participants were asked to complete 200 no reward trials. Our results show that even after 24 hours, and over the course of 200 additional unrewarded trials, participants maintained similar MT performance levels. We used a repeated measures ANOVA with timepoint (early vs late across all testing days) as the within factor to assess changes across testing days (repeated measures ANOVA, main effect for timepoint, F = 28.65, p < 0.0001; Figure 8a; Supplementary Figure 4). These results indicate that performance improved over the course of the experiment. However, no changes in MT performance could be observed between late training on day 2 and early training on day 3 (Wilcoxon test, Z = −1.21, p = 0.3016) and between early and late training on day 3 (Wilcoxon test, Z = −1.48, p = 0.4444). Similarly, coarticulation levels appeared stable across the additional testing day without reward (repeated measures ANOVA, main effect for timepoint, F = 19.19 p < 0.0001; Figure 8b; Supplementary Figure 4), with no changes in performance between late day 2 and early day 3 (Wilcoxon test, Z = −0.94, p = 1). Similarly to MT, no changes were observed across day 3 (Wilcoxon test; early vs late; day 3, Z = −0.67, p = 1). Yet again, we found that peak velocities were persevered transitioning to and across day 3 (Wilcoxon test, late training day 2 x early training day 3, Z = −1.21, p = 1; early training day 3 x late training day 3, Z = −1.21, p = 1; Figure 8c; Supplementary Figure 4). Taken together, these findings highlight that performance gains were preserved even after an additional testing day without reward availability.

**Figure 8.**
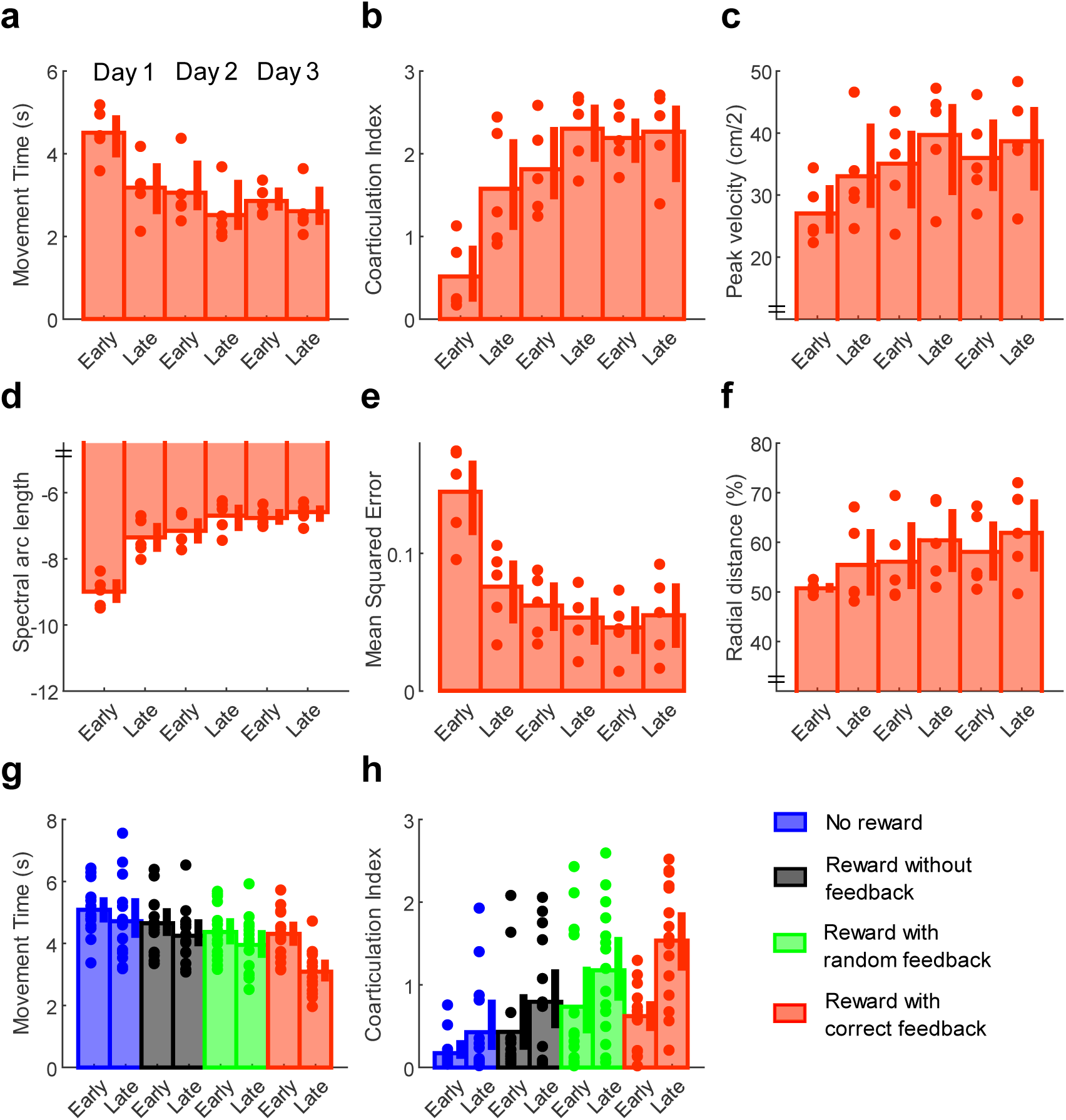
Long-term maintenance of performance without reward. a-f: Experiment 2. Median data across all 3 days (early (first 20 trials) vs late (last 20 trials) training). **a)** MT **b)** CI level **c)** Peak velocity **d)** Spectral arc length (smoothness) **e)** Mean squared error between data and minimum jerk model prediction **f)** Radial distance (spatial reorganisation) g**-h: Experiment 3. g)** Median MT across groups. **h)** Median CI level.

When assessing changes in smoothness using spectral arc length, we found similar results to experiment 1. Smoothness improved over the course of the experiment (repeated measures ANOVA, main effect for timepoint, F = 57.14 p < 0.0001; Figure 8d; Supplementary Figure 4), while no changes could be observed between late training on day 2 and early training on day 3 (Wilcoxon test, Z = −0.67, p = 1) and between early and late training on day 3 (Wilcoxon test, Z = −1.21, p = 1). In line with these results, we found that performance aligned progressively with the predictions of the minimum jerk model (repeated measures ANOVA, main effect for timepoint, F = 23.38 p < 0.0001; Figure 8e; Supplementary Figure 4), while no significant changes in similarity could be observed transitioning to and across day 3 (Wilcoxon test, late training day 2 x early training day 3, Z = −2.02, p = 1; early training day 3 x late training day 3, Z = −1.21, p = 1). In addition, we found that participants progressively reorganised their spatial movement output, with a significant decrease in radial distance found across the experiment (repeated measures ANOVA, main effect for timepoint, F = 57.14 p < 0.0001; Figure 8f; Supplementary Figure 4). However, no changes in radial distance could be observed between late training on day 2 and early training on day 3 (Wilcoxon test, Z = - 2.02, p = 1) and between early and late training on day 3 (Wilcoxon test, Z = −1.75, p = 1). These findings support our results from experiment 1 and indicate that improvements in movement efficiency through a quantitative change in how the task is performed may enable long-term retention of reward-based performance gains.

### Reward based on performance is most effective in producing behavioural change

Our third experiment was intended to assess whether the observed changes on day 1 were specific to training with reward-based feedback of performance. Participants were allocated to one of the four groups: (1) no reward, (2) reward without performance-based feedback, (3) reward with random feedback and (4) reward with accurate feedback. Only groups 2-4 received monetary reward; reward for groups 1 and 4 were equivalent to that used in experiment 1. Participants in group 3 received feedback about the reward delivered on each trial which was randomly drawn from the feedback experienced during experiment 1. It therefore matched group 4 in terms of reward probability but this did not correspond to actual performance. In contrast, participants in group 2 did not receive any performance-based feedback after completing a given trial. Upcoming reward trials, however, were still cued. Since groups 1 and 4 underwent the same regime as the no reward and reward group in experiment 1 respectively, we were able to test whether our results replicated. Group 4 produced the largest improvements in performance, being the only group who showed significant improvements in MT (mixed ANOVA; timepoint x group; interaction, F = 3.76, p = 0.0158; Wilcoxon test; early vs late group 4: Z = 3.40, p = 0.0028; Figure 8g; Supplementary Figure 5) and coarticulation (mixed ANOVA; timepoint x group; interaction, F = 3.18, p = 0.0308; Wilcoxon test; early vs late group 4: Z = −3.24, p = 0.0048; Figure 8h; Supplementary Figure 5). This indicates that the expectation of reward is not sufficient to induce large behavioural improvements with a link to performance being essential to optimise gains.

These results showed that performance-based reward invigorates sequential reaching. Driven by a reward-based increase in speed, movements also exhibited greater coarticulation, smoothness and a closer alignment to the predictions of a minimum jerk model. Importantly, these performance gains were maintained across multiple days even after the subsequent withdrawal of reward. This highlights the importance of coarticulation to skilful sequential reaching performance, and the potential of this mechanism to produce long-lasting reward-driven improvements in behaviour.

## Discussion

Motor skill learning is integral to everyday life and describes improvements in performance above baseline levels ^59^. Improvements in skill have often been assessed at the levels of speed, accuracy and more recently efficiency ^60^. In our task, improvements in speed (MT) could be achieved via two strategies: (1) increases in peak velocities of the individual reaching movements and (2) reduction in dwell times around the via points (coarticulation). We were able to show that reward invigorated peak velocities, supporting previous findings on the reward-based invigoration of simple discrete reaching movements ^41,42^. Moreover, for the first time, we demonstrate that reward accelerates the slow learning process associated with coarticulation ^9–11,31^. With respect to movement efficiency, these two strategies differ regarding the metabolic costs they incur. Recently, it has been demonstrated that in discrete reaching movements, improvements in peak velocity can be achieved at no cost to accuracy through increased arm stiffness ^41^. Although an attractively simple strategy, it comes with a marked escalation in metabolic costs ^56^. In fact, it has been shown that in discrete reaching tasks such invigoration is reward-dependent and transient in nature ^40–42^. We suggest that this could be due to the energetic demands of the invigorated movement requiring the continued presence of reward to negate this added cost. In contrast, coarticulation enables the performance of sequential movements to become similar to a minimum jerk trajectory through the merging of neighbouring movements into singular smooth elements ^9–12,53^. Increases in smoothness have been shown to reduce metabolic costs, thereby enhancing overall movement efficiency ^54^. Here we showed that performance in the reward group was smoother and exhibited greater similarity to a minimum jerk trajectory. The subsequent hypothesised improvement in energetic efficiency might explain why the enhanced performance of the reward group became reward-independent, whereas performance in the no reward group, who showed little coarticulation, followed the previously observed transient ‘on-off’ effect.

Despite previous studies demonstrating that reward can enhance retention across a wide variety of sequential and continuous motor tasks, the underlying mechanism for this effect has been described at a very abstract level ^50–52^. Specifically, it is unclear how reward strengthens a motor memory so that improved performance is maintained even when the incentive is no longer provided. We believe that for reward to induce such long-term improvements in motor skill, it must not only lead to enhanced performance but also improvements in efficiency. Reduction in the cost of this enhanced performance (faster or more accurate movements ^40^), enables it to be performed long-term without further incentive/reward. Therefore, reward may not enhance the memory of the action but instead lead to the task being performed in a fundamentally different, and more efficient, manner. In support of this, we found that participants in the reward group produced a quantitative change (spatial reorganisation of the velocity profile between neighbouring movements) in how the task was executed. This highlights that careful analysis of how reward influences the performance of complex actions is essential for dissociating its effects on execution and retention.

Although coarticulation might appear to feature similar movement characteristics as chunking, it is important to emphasise that they represent different processes. Chunking often refers to a series of discrete movements (e.g. button presses in sequence learning paradigms) which are temporally aligned (chunked) over the course of training ^7,26–29^. The chunks are represented at a behavioural level through shorter reaction times between actions, and often allow for a faster execution overall ^7,8^. Yet the elements within a chunk are still performed discretely with a clear stop period between them ^22,29^. In addition, at a neural level, these elemental movements are planned individually through competitive queuing, highlighting again that some level of independence is maintained ^61^. In contrast, coarticulation reflects the merging of neighbouring movements into a single motor primitive that no longer features a pronounced stop period and must run to completion once initiated ^9–11,31^. While kinematically appearing to represent a single movement, it remains an open question how these newly formed primitives are represented at a neural level in humans.

Recent work in animals suggests that sequence learning is a dopamine-dependent process ^62,63^. By tracking dopamine concentration changes in the nucleus accumbens core of rats, it has been shown that dopamine levels dynamically change with task proficiency ^62^. Further evidence originates from research in parkinson’s disease (PD)in which PD patients OFF medication exhibit specific impairments in movement chunking ^32,33^. However, whether dopamine underpins coarticulation similarly to chunking is still unexplored (a notable exception being ^33^). Specifically, whether dopamine directly acts on coarticulation or whether any mediating relationship is due to the effects of dopamine on movement invigoration, requires further examination ^33^.

Here, we show that coarticulation is associated with smoother and, with regards to minimising jerk, more efficient execution. Interestingly, reaching movements performed by stroke patients exhibit reduced smoothness ^34–37^, with increases in jerk being due to a decomposition of a movement into a series of sub-movements ^34–37^. However, over the course of the recovery process, performance becomes smoother as these sub-movements are progressively blended ^34–37^. Considering this theoretical proximity to the concept of coarticulation, we speculate that stroke recovery and coarticulation may follow similar principles. Consequently, coarticulation facilitated by reward could be a powerful tool in stroke rehabilitation to promote smooth and efficient sequential actions which form an essential component of everyday life activities.

In conclusion, this work highlights that coarticulation provides a mechanism by which reward can invigorate sequential performance whilst also improving efficiency. This improvement in efficiency through a quantitative change in how the action is performed appears essential for the retention of reward-based improvements in motor behaviour.

## Methods

### Participants

107 participants (17 males; age range 18 - 35) were recruited to participate in three experiments, which had been approved by the local research ethics committee of the University of Birmingham. All participants were novices to the task paradigm and were free of motor, visual and cognitive impairment. Most participants were self-reportedly right-handed (N = 7 left-handed participants) and gave written informed consent prior to the start of the experiment. For their participation, participants were remunerated with either course credits or money (£7.5/hour) and were able to earn additional money during the task depending on their performance. Depending on the experiment, participants were pseudo-randomly allocated to one of the available groups.

### Experimental Apparatus

All experiments were performed using a Polhemus 3SPACE Fastrak tracking device (Colchester, Vermont U.S.A; with a sampling rate of 110Hz). Participants were seated in front of the experimental apparatus which included a table, a horizontally placed mirror 25cm above the table and a screen (Figure 1a). The low-latency Apple Cinema placed 25cm above the mirror had a refresh rate of 60Hz and displayed the workspace and paricipants’ hand position (represented by green cursor – diameter 1cm). On the table, participants were asked to perform 2-D reaching movements. Looking into the mirror, they were able to see the representation of their hand position reflected from the screen above. This setup effectively blocked their hand from sight. The experiment was run using MATLAB (The Mathworks, Natwick, MA), with Psychophysics Toolbox 3.

### Task design

Participants were asked to hit a series of targets displayed on the screen (Figure 1c). Four circular (1cm diameter) targets were arranged around a centre targe (‘via target‘). Starting in the via target, participants had to perform eight continuous reaching movements to complete a trial. Target 1 and 4 were displaced by 10cm on the *y-*axis, whereas Target 2 and 3 were 5cm away from the via target with an angle of 126 degrees between them (Figure 1c). Our task design was based on previous work ^9–11,31^ in which the authors were able to observe coarticulation using similar angles and reaching distance configurations. To start each trial, participants had to pass their cursor though the preparation box (2×2cm) on the left side of the workspace, which triggered the appearance of the start box (2×2cm) in the centre of the screen. After moving the cursor into the start box, participants had to wait for 1.5s for the targets to appear. This ensured that participants were stationary before reaching for the first target. Target appearance served as the go-signal and the start box turned into the via target (circle). Upon reaching the last target (via target), all targets disappeared, and participants had to wait for 1.5s before being allowed to exit the start box to reach for the preparation box to initiate a new trial. Participants had to repeat a trial if they missed a target or performed the reaching order incorrectly. Similarly, exiting the start box too early either at the beginning or at the end of each trial resulted in a missed trial.

### Reward Structure and Feedback

Participants in experiment 1 and experiment 2 experienced either reward or no reward trials depending on the current experimental phase: (1) Reward trials were cued using a visual stimulus prior to the start of the trial. Once participants moved into the preparation box, the start box appeared in yellow (visual stimulus) rather than in black (Figure 1f). Participants were informed that faster MTs will earn them more money, with a maximum amount of 5p available in each trial. While participants moved from the start box to the preparation box to initiate a new trial, the amount earned in the previous trial was displayed on the top of the screen (i.e ‘you have earned 2p out of 5p‘). We used a closed-loop design to calculate the amount of reward earned in each trial. To calculate this, we included the MT values of the last 20 trials and organised them from fastest to slowest to determine the rank of the current trial within the given array. A rank in the top three (<= 90%) returned a value of 5p, ranks >= 80% and <90% were valued at 4p; ranks >=60% and <80% were awarded 3p; ranks >=40% and < 60% earned 2p while 1p was awarded for ranks >=20% and < 40%. A rank in the bottom three (<20%) returned a value of 0p. When participants started a new experimental block, performance in the first trial was compared to the last 20 trials of the previously completed block. (2) No reward trials were not cued, and no reward was available for participants. However, participants were instructed to “move as fast and accurately as possible”. experiment 3, participants were randomly allocated to one of the four groups: (1) no reward, (2) reward without performance-based feedback, (3) reward with random feedback and (4) reward with accurate feedback. Participants in the no reward (1) and reward with accurate feedback (4) groups underwent the same regime as the no reward and reward groups in experiment 1 respectively. To investigate whether reward and/or feedback drove performance changes, we changed the reward and feedback structure for the two remaining groups. Participants in the reward without performance-based feedback group (2) were able to earn money depending on their performance in each trial. Reward trials were cued with a yellow start box prior to the start of the trial. However, participants did not receive any feedback on their performance after completing a given trial. They were asked to initiate a new trial and ‘be as fast and accurate as possible to earn more money’. In contrast to this, participants in the reward with random feedback group (3) received feedback after completing a reward trial during training, which were also cued with a yellow start box. However, feedback in this group was not performance-based, but was drawn randomly from feedback given to participants in experiment 1. To this end, we strung together all reward values given to participants in the first experiment and randomly chose a value for feedback in a given trial in experiment 3. Participants, therefore, received feedback which was similar in reward probability without corresponding to actual performance.

## Experimental Procedure

### Experiment 1

In this experiment, we investigated whether reward can invigorate performance on a sequential reaching task. The experiment included an initial learning phase prior to the start of the experiment as well as a baseline, training and two post assessments. The same design without the learning phase was repeated 24 hours later (Figure 1b). Participants were pseudo-randomly allocated to either the reward or no reward group (N = 21 each) and were informed that at some point during the experiment they would be able to earn additional money depending on their performance.

#### Learning

We included a learning phase prior to the start of the experiment for participants to be able to memorise the reaching sequence. This allowed us to attribute any performance gains to improvements in execution rather than memory. Once participants waited 1.5s inside the start box, the targets appeared which were numbered clockwise from 1 to 4 starting with the central top target. Participants were also able to see a number sequence at the top left of the screen displaying the order of target reaches (1 – 3 – 2 - 4). Participants were instructed to hit the targets according to the number sequence while also hitting the via target in between target reaches. They had to repeat a trial if they missed a target or performed the reaching order incorrectly. Similarly, exiting the start box too early either at the beginning or at the end of each trial resulted in a missed trial. After a cued trial, participants were asked to complete a trial from memory without the number sequence or numbers inside the targets. If participants failed a no cue trial more than twice, cues appeared in the following trial as a reminder. After a maximum of 10 cue and 10 no cue trials participants completed this block.

#### Baseline

Participants in both groups completed 10 baseline trials, which were used to assess whether there were any pre-training differences between groups Both groups were instructed to ‘move as fast and accurately as possible“, while no performance-based feedback was given at the end of each trial.

#### Training

Participants in the reward group were informed that during this part they would be able to earn money depending on how fast they complete each trial (200 reward trials). In contrast, participants in the no reward group engaged in 200 no reward trials and were again instructed to move as fast and as accurately as possible.

#### Post assessments

On both testing days, participants from both groups were asked to complete two post assessments (20 trials each); one with reward trials (post-R) and one with no reward trials (post-NR). The order was counter-balanced across participants.

### Experiment 2

In this experiment, we aimed to test how robust reward-driven performance gains were over an additional testing day without reward availability. Participants (N = 5) underwent the same regime as the reward group in experiment 1 on the first two testing days. On the third testing day after baseline, participants were asked to complete 200 no reward trials.

### Experiment 3

Here, we investigated whether the observed performance gains depend on reward expectation, performance-based feedback or a combination of both. To this end, we allocated participants (N = 60) to one of the four groups: (1) no reward, (2) reward without performance-based feedback, (3) reward with random feedback and (4) reward with accurate feedback (see Reward Structure and Feedback for more information). Participants underwent the same procedure as participants in experiment 1 over the course of the first testing day. After a learning phase and a baseline part, participants engaged in 200 training trials which differed with regards to their reward and feedback structure. Similarly, to experiment 1 and 2, participants then completed the post assessments, which were counter-balanced across participants.

## Data Analysis

Analysis code is available on the Open Science Framework website, alongside the experimental datasets at https://osf.io/62wcz/. The analyses were performed in Matlab (Mathworks, Natick, MA).

## Movement Time (MT)

MT was measured as the time between exiting the start box and reaching the last target. This excludes reaction time, which describes the time between target appearance and when the participants’ start position exceeded 2cm. Trials with MTs beyond 9.0s were excluded from analysis, which amounted to 0.37% of all trials.

### Peak Velocity

Through the derivative of positional data (*x, y*), we obtained velocity profiles for each trial which were smoothed using a gaussian smoothing kernel (σ = 2). The velocity profile was then divided into segments representing movements to each individual target (8 segments) by identifying when the positional data was within 2cm of a target. We measured the peak velocity (*v*_*peak*_) of each segment by finding the maximum velocity:

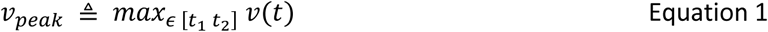

Where *v(t)* is the velocity of segment *t*, and *t*_*1*_ *and t*_*2*_ represent the start and end of segment *t* respectively.

### Coarticulation Index (CI)

Coarticulation describes the blending together of individual motor elements into a singular smooth action. This is represented in the velocity profile by the stop period between the two movements gradually disappearing and being replaced by a single velocity peak (Figure 1c) ^9–11^. Based on the predictions of a model which minimised jerk (Equation 3; Figure 1d), we hypothesised that coarticulation would mainly occur between out-centre centre-out reaching movements (Figure 1e). We therefore excluded the first and last target reach from this analysis. To measure coarticulation, we compared the mean peak velocities of two sequential reaches with the minimum velocity around the via point. The smaller the difference between these values, the greater coarticulation had occurred between the two movements (Figure 2c) ^25^. The velocity profile was cut into 3 segments depending on the peak velocities of the out-centre and centre-out reaching movements. The minimum velocity of these segments was calculated and compared to the average of the peak velocities:

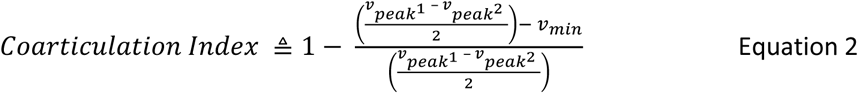

with *v*_*peak1*_ and *v*_*peak2*_ representing the velocity peak value of the out-centre and centre-in reaching movement of a given segment, respectively, and *v*_*min*_ representing the minimum value between these two points. We normalised the obtained difference, ranging from 0 to 1, with 1 indicating a fully coarticulated movement. Given that in this task three pairs of movements were able to be coarticulated, the maximum CI value was 3 in each trial. Note that this measure is MT insensitive since it focuses on the difference in values not on their absolute value. This means slow and fast movements could have a similar CI value.

### Error

We operationalised error as the amount of repetitions necessary to complete a given trial. Trials had to be repeated if participants missed a target or if they exited the start box before the targets appeared or disappeared.

### Minimum-jerk model

A traditional minimum-jerk model for motor control is guided by optimisation theory, where a cost is minimised over the trajectory^12,53^. In the case of the minimum-jerk model, the cost is defined as the squared jerk (3^rd^ derivative of position with respect to time):

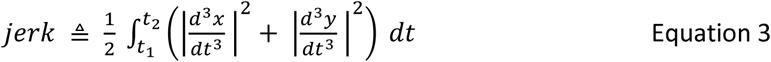

Here *x* and *y* represent the position of the index finger over time (*t*), while *t*_*1*_ and *t*_*2*_ define the start and end of a trial in seconds (*t*). The Matlab code provided by Todorov and Jordan (1998)^64^ was used to compute the minimum jerk trajectory (trajectory that minimised Equation 3), and the accompanying velocity profile, given a set of via points, start/end position and movement time ^12^. We then calculated the mean square error (immse function in Matlab) between the predicted and actual velocity profile, which were both normalised and interpolated (N = 500), to estimate the fit on a trial-by-trial basis. Due to the two-dimensional structure of trajectories, we chose the velocity profiles of rather than the trajectories for this comparison.

### Spectral Arc Length

To further assess movement smoothness, we measured spectral arc length. Although we decided to use the traditional jerk metric in our modelling analysis, to allow for comparisons with prior literature, spectral arc length has been shown to be less sensitive to differences in MT and more sensitive to changes in smoothness ^55,65^. The spectral arc length is derived from the arc length of the power spectrum of a Fourier transformation of the velocity profile. We used an open-source Matlab toolbox to calculate this value for each trajectory ^66^.

For both spectral arc length and the minimum-jerk model, we only included non-corrected trials. Trials that were classified as corrected included at least one corrective movement to hit a previously missed target. These additional movements added peaks to the velocity profile which complicated model comparison and increased jerkiness disproportionally. Therefore, 1820 trials were excluded for both analyses (8.68% of all trials).

### Spatial Reorganisation

In addition to CI, coarticulation can also be expressed spatially as the radial distance between the peak velocity (v_peak_) on the sub-movements and the minimum velocity (v_min_) around the via point (Figure 7a). This distance becomes smaller with increased coarticulation ^9,10^ and reflects the merging of two sub-movements into one (Supplementary Figure 3). To measure these changes in radial distance between peaks and via points, we used a sliding window approach of 10 trials at a time. For each target reach (excluding the first and the last) we fitted a confidence ellipse ^67^ with a 95% confidence criterion around the scatter of the spatial position (*x, y*) of each peak velocity of the included trials (Figure 7a). The confidence ellipses were obtained using principal component analysis to determine the minimum and maximum dispersion of the included data points in the *x-y* plane. To measure the distance between the scatter and its corresponding via point, we determined the ellipse’s centoid (point of intersection of ellipse’s axes) and calculated the radial distance to the via point. The obtained distance values were normalised and ranged from 0-100%, with 100% representing 0 cm distance between the centroid and the via point. Considering that individual reaching movements display a bell-shaped velocity profile, with the *v*_*peak*_ situated approximately in the centre of the movement, radial distance values between 45-55% can be expected if each movement is executed individually (Supplementary Figure 3). To understand the optimal radial distance to the via point, we measured the radial distance for a trajectory which minimised jerk. This suggested that values of 82-85% represent the optimal range for coarticulated movements in this task (Figure 7g).

### Variability

To assess changes in variability we measured the area of the peak velocity ellipses using the same approach as for spatial reorganisation. The area of the ellipse represents the total variance of the included data points in the *x-y* plane and was calculated by multiplying the axes of the ellipse with pi (π) ^67,68^. Data was normalised to the baseline for each group.

## Statistical Analysis

Wilcoxon tests were used to analyse differences in performance during baseline. Mixed model ANOVAs were used to assess statistical significance of our results in experiment 1. We carried out separate analyses for training with timepoint (early training (first 20 trials), late training (last 20 trials)) and group (reward, no reward), and post assessment with condition (post-R, post-NR (all 20 trials in each)) and group (reward, no reward) as factors for both days. We used one-sample Kolmogorov-Smirnov tests to test our data for normality and found that all measures were non-parametric. Median values were therefore used as input in all mixed model ANOVAs. Wilcoxon tests were employed when a significant interaction and/or main effects were reported and corrections for multiple comparisons were performed using Bonferroni correction. Linear partial correlations (fitlm function in Matlab) were used to measure the degree of association between the chosen variables, while accounting for the factor group. Piecewise linear spline functions were fitted through the scatter of spatial distance values and CI levels using least square optimisation by means of shape language modelling (SLM) ^69^. We used three knots as input for the linear model.

A repeated-measure ANOVA was used to test for significance of our results in experiment 2. We compared MT and CI performance separately with timepoint (early training, late training over all 3 testing days) as the within factor. Due to our data being non-parametric after using one-sample Kolmogorov-Smirnov tests, we included median values as input for all repeated-measure ANOVAs. Wilcoxon test was used as post-hoc test and multiple comparisons were corrected using Bonferroni corrections.

We used mixed model ANOVAs to statistically analyse our results from experiment 3. Separate analyses were carried out for training and post assessment for both MT and CI. Timepoint (early training, late training) and group (no reward, reward without feedback, reward with random feedback, reward with correct feedback) were factors to assess performance in training. Post assessment performance was analysed with a mixed model ANOVA that had condition (Post-R, Post-NR) and group (no reward, reward without feedback, reward with random feedback, reward with correct feedback) as factors. We used median values as input, because one-sample Kolmogorov-Smirnov tests confirmed a lack of normality. Wilcoxon tests were employed when a significant interaction and/or main effects were reported and corrections for multiple comparisons were performed using Bonferroni correction.

## Acknowledgements

This work was supported by the European Research Council starting grant: MotMotLearn (637488)

## Supplement

**Supplementary Figure 1.**
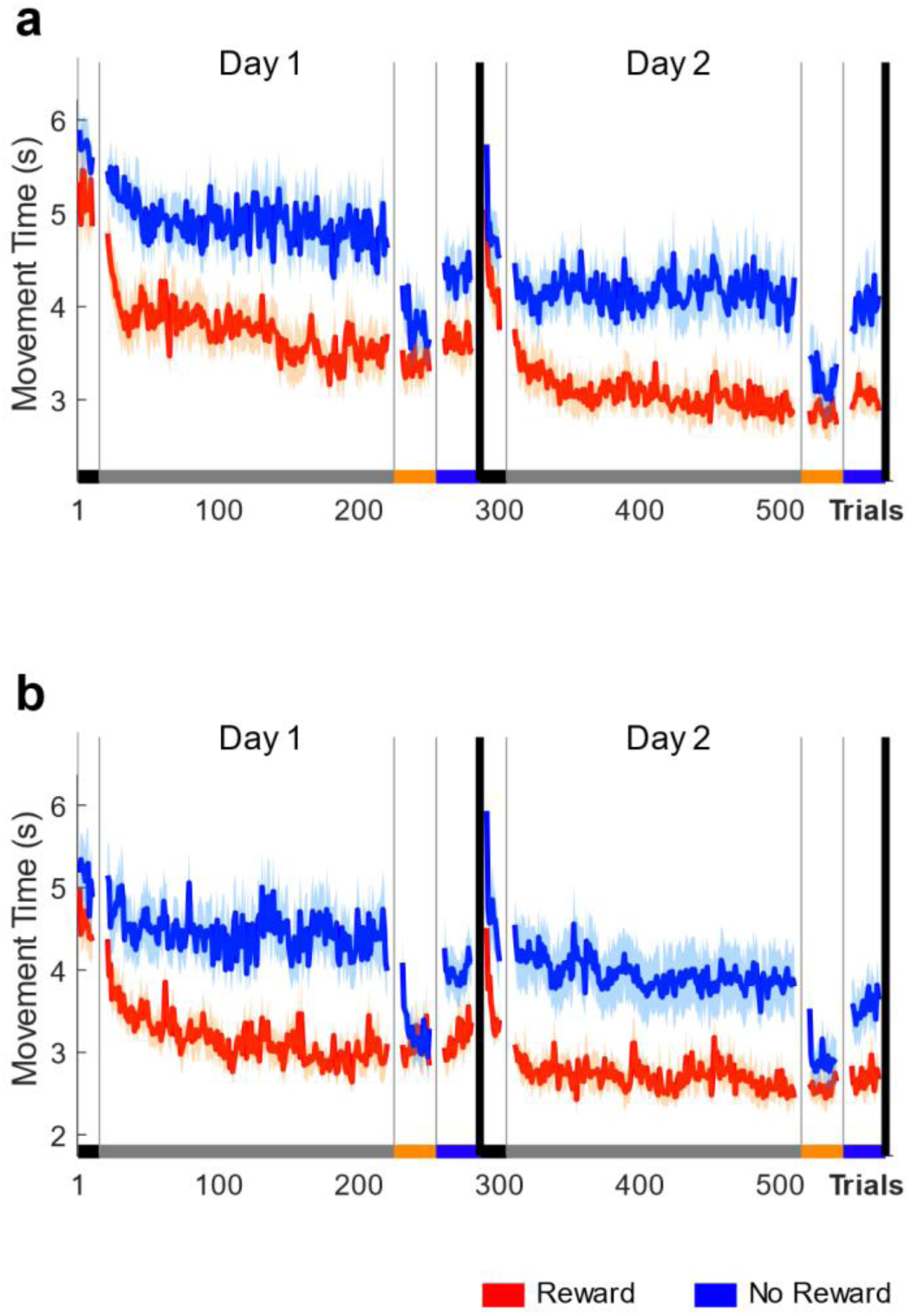
Counterbalancing did not affect MT results. Data was split depending on the counterbalancing. Participants in **a)** completed post-R prior to post-NR, whereas participants encountered the reversed order during their post assessment (**b**).

**Supplementary Figure 2.**
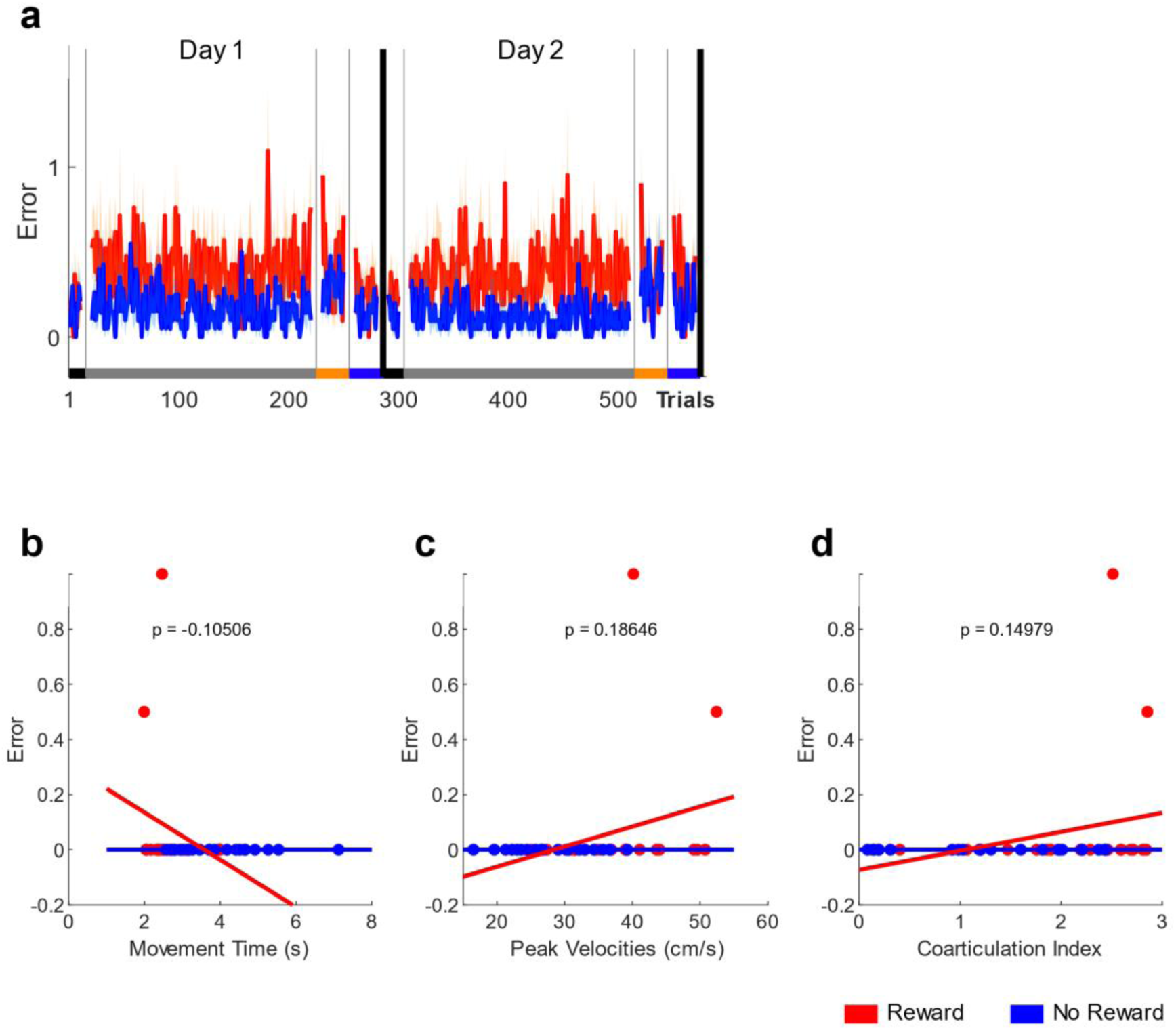
Number of errors did not affect performance during later stages of training. **a)** Trial-by-trial data showing the number of errors across participants for both groups). **b-d)** Scatterplots comparing median error data during early training with median **(b)** MT, **(c)** peak velocity and **(d)** CI data during late training.

**Supplementary Figure 3.**
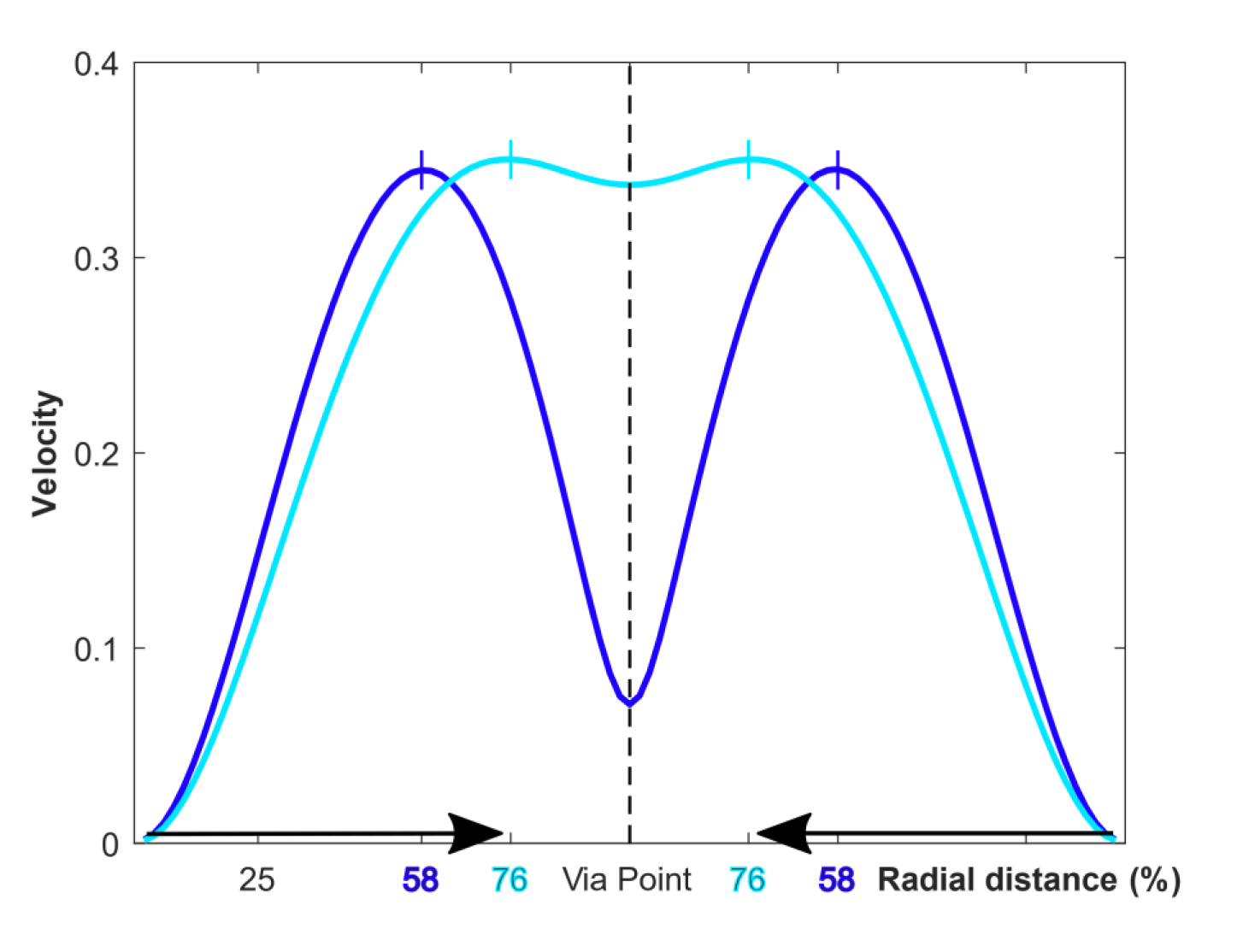
Illustration of radial distance (%) values depending on different levels of coarticulation. Considering two target reaches (same length) the spatial position (*x,y*) changes depending on the level of coarticulation thereby exhibiting less radial distance to the via point. We used the predictions of the minimum jerk model used in this paper to plot the optimal solutions for the here included sequential target reaches.

**Supplementary Figure 4.**
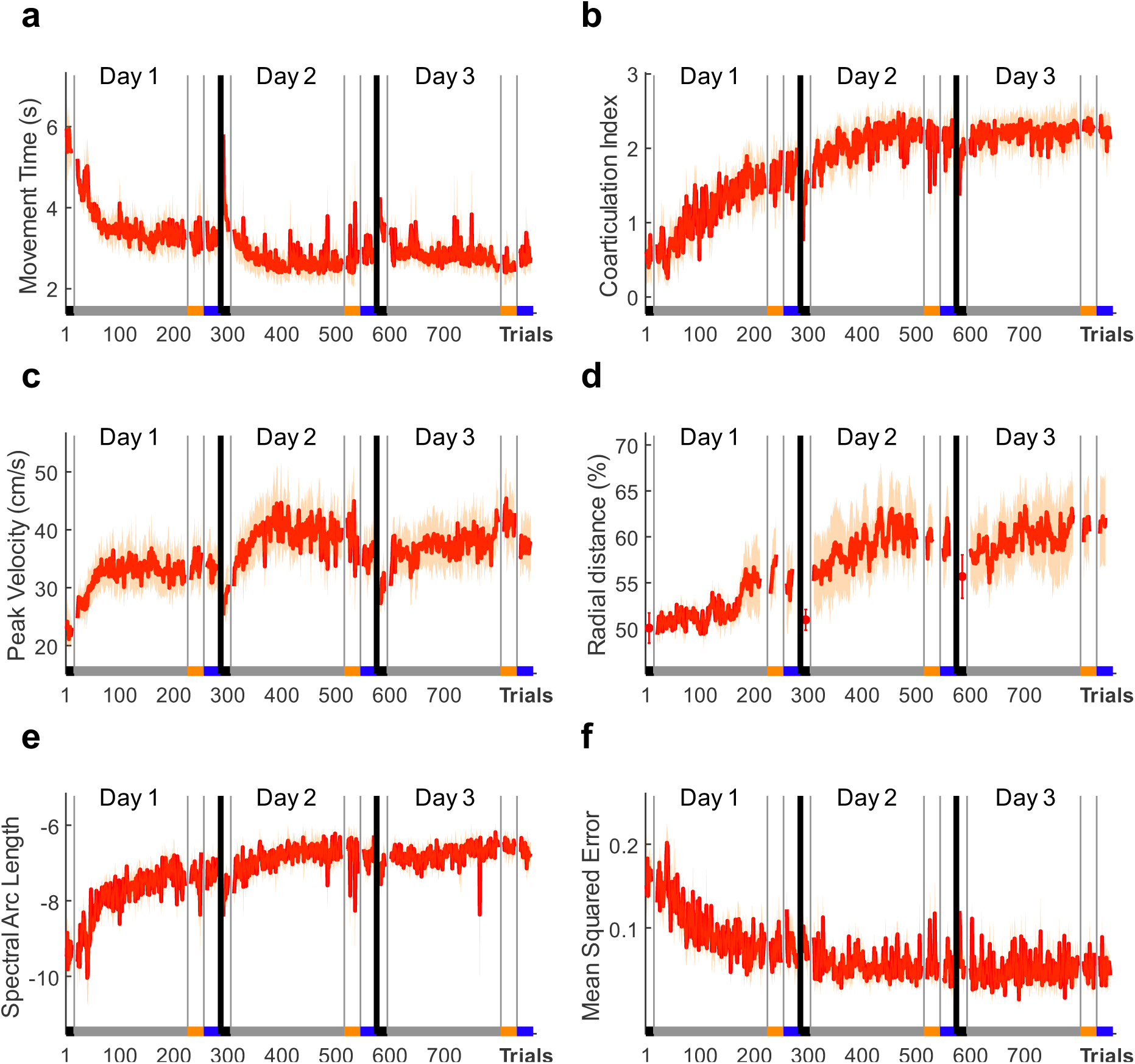
Performance gains are stable even over a prolonged washout period. Trial-by-trial data for **a)** MT, **b)** CI levels **c)** peak velocity **d)** radial distance, **e)** spectral arc length, **f)** mean squared error CI for the second experiment which was scheduled on three consecutive days to assess behavioural change over a prolonged washout period. Participants received reward during training on the first two days, however no reward was available on the third day.

**Supplementary Figure 5.**
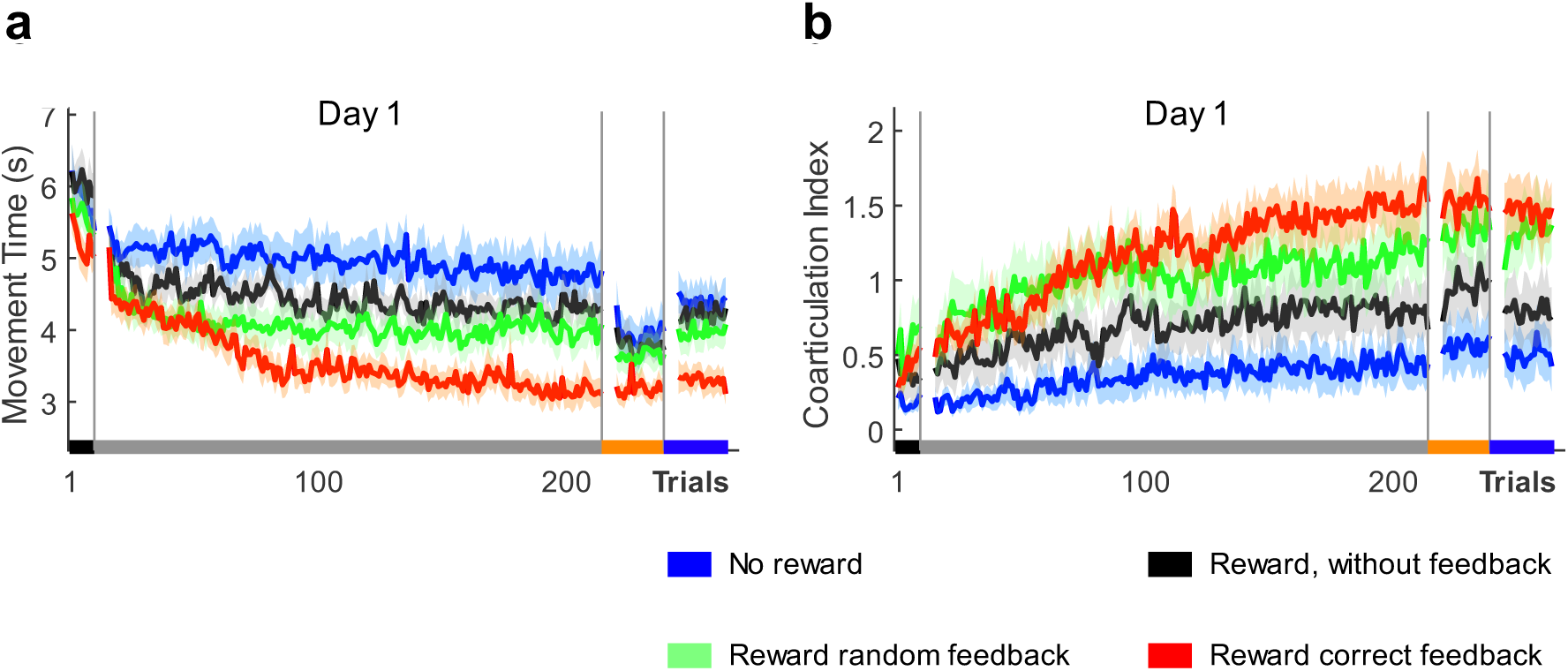
Reward based on correct performance feedback is most efficient at driving behavioural change. Trial-by-trial data for **a)** MT and **b)** CI levels for the third experiment which included 4 groups that differed with regards to reward availability and feedback type during training to investigate what drove behavioural changes observed in experiment 1.

## References

1. Fowler, C. A. Coarticulation and theories of extrinsic timing. 21.

2. Willingham, D. B. A Neuropsychological Theory of Motor Skill Learning. 27.

3. Doeringer, J. A. & Hogan, N. Serial processing in human movement production. Neural Networks 11, 1345–1356 (1998).

4. Jin, X., Tecuapetla, F. & Costa, R. M. Basal ganglia subcircuits distinctively encode the parsing and concatenation of action sequences. Nat Neurosci 17, 423–430 (2014).

5. Shah, A., Barto, A. & Fagg, A. A Dual Process Account of Coarticulation in Motor Skill Acquisition. Journal of Motor Behavior 45, 531 (2013).

6. Hansen, E., Grimme, B., Reimann, H. & Schöner, G. Anticipatory coarticulation in non-speeded arm movements can be motor-equivalent, carry-over coarticulation always is. Exp Brain Res 236, 1293–1307 (2018).

7. Diedrichsen, J. & Kornysheva, K. Motor skill learning between selection and execution. Trends Cogn. Sci. (Regul. Ed.) 19, 227–233 (2015).

8. Ramkumar, P. et al. Chunking as the result of an efficiency computation trade-off. Nat Commun 7, 12176 (2016).

9. Sosnik, R., Hauptmann, B., Karni, A. & Flash, T. When practice leads to co-articulation: the evolution of geometrically defined movement primitives. Exp Brain Res 156, 422–438 (2004).

10. Sosnik, R., Chaim, E. & Flash, T. Stopping is not an option: the evolution of unstoppable motion elements (primitives). Journal of Neurophysiology 114, 846 (2015).

11. Sosnik, R., Flash, T., Hauptmann, B. & Karni, A. The acquisition and implementation of the smoothness maximization motion strategy is dependent on spatial accuracy demands. Exp Brain Res 176, 311–331 (2007).

12. Todorov, E. & Jordan, M. I. Smoothness Maximization Along a Predefined Path Accurately Predicts the Speed Profiles of Complex Arm Movements. Journal of Neurophysiology 80, 696–714 (1998).

13. Kuhnert, B. & Nolan, F. The origin of coarticulation. FIPKM) 35, (1997).

14. Kent, R. D. & Minifie, F. D. Coarticulation in recent speech production models. Journal of Phonetics 5, 115–133 (1977).

15. Jerde, T. E., Soechting, J. F. & Flanders, M. Coarticulation in Fluent Fingerspelling. The Journal of Neuroscience 23, 2383 (2003).

16. Winges, S. A. & Furuya, S. Distinct digit kinematics by professional and amateur pianists. Neuroscience 0, 643 (2015).

17. Soechting, J. F. & Flanders, M. Organization of sequential typing movements. Journal of Neurophysiology (1992) doi: 10.1152/jn.1992.67.5.1275.

18. Soechting, J. F. & Flanders, M. Flexibility and Repeatability of Finger Movements During Typing: Analysis of Multiple Degrees of Freedom. J Comput Neurosci 4, 29–46 (1997).

19. Rumelhart, D. E. & Norman, D. A. Simulating a Skilled Typist: A Study of Skilled Cognitive-Motor Performance. Cognitive Science 6, 1–36 (1982).

20. Breteler, M. D. K., Hondzinski, J. M. & Flanders, M. Drawing Sequences of Segments in 3D: Kinetic Influences on Arm Configuration. Journal of Neurophysiology (2003) doi: 10.1152/jn.01062.2002.

21. Verwey, W. B. & Dronkert, Y. Practicing a Structured Continuous Key-Pressing Task: Motor Chunking or Rhythm Consolidation? J Mot Behav 28, 71–79 (1996).

22. Verwey, W. B., Groen, E. C. & Wright, D. L. The stuff that motor chunks are made of: Spatial instead of motor representations? Exp Brain Res 234, 353–366 (2016).

23. Wymbs, N. F., Bassett, D. S., Mucha, P. J., Porter, M. A. & Grafton, S. T. Differential recruitment of the sensorimotor putamen and frontoparietal cortex during motor chunking in humans. Neuron 74, 936 (2012).

24. Thompson, J. J., McColeman, C. M., Blair, M. R. & Henrey, A. J. Classic motor chunking theory fails to account for behavioural diversity and speed in a complex naturalistic task. PLOS ONE 14, e0218251 (2019).

25. Friedman, J. & Korman, M. Observation of an expert model induces a skilled movement coordination pattern in a single session of intermittent practice. Sci Rep 9, 1–15 (2019).

26. Boyd, L. A. et al. Motor sequence chunking is impaired by basal ganglia stroke. Neurobiology of learning and memory 92, 35–44 (2009).

27. Huntley, J., Bor, D., Hampshire, A., Owen, A. & Howard, R. Working memory task performance and chu k y A zh m ‘ Br J Psychiatry 198, 398–403 (2011).

28. Ariani, G. & Diedrichsen, J. Sequence learning is driven by improvements in motor planning. J. Neurophysiol. 121, 2088–2100 (2019).

29. Verwey, W. B. Evidence for a multistage model of practice in a sequential movement task. J EXP PSYCHOL HUMAN 25, 1693–1708 (1999).

30. Barnhoorn, J. S., Van Asseldonk, E. H. F. & Verwey, W. B. Differences in chunking behavior between young and older adults diminish with extended practice. Psychological Research 83, 275–285 (2019).

31. Sosnik, R., Flash, T., Sterkin, A., Hauptmann, B. & Karni, A. The activity in the contralateral primary motor cortex, dorsal premotor and supplementary motor area is modulated by performance gains. Frontiers in Human Neuroscience 8, (2014).

32. Bidet-Ildei, C., Pollak, P., Kandel, S., Fraix, V. & Orliaguet, J.-P. Handwriting in patients with Parkinson disease: effect of L-dopa and stimulation of the sub-thalamic nucleus on motor anticipation. Hum Mov Sci 30, 783–791 (2011).

33. M zz,, H v, A. & K k u, J. W. Why D’ W M v F ? k’ D, Movement Vigor, and Implicit Motivation. J. Neurosci. 27, 7105–7116 (2007).

34. Rohrer, B. et al. Movement Smoothness Changes during Stroke Recovery. The Journal of Neuroscience 22, 8297 (2002).

35. Rohrer, B. et al. Submovements Grow Larger, Fewer, and More Blended during Stroke Recovery. Motor Control 8, 472–483 (2004).

36. Dipietro, L., Krebs, H. I., Fasoli, S. E., Volpe, B. T. & Hogan, N. Submovement changes characterize generalization of motor recovery after stroke. Cortex 45, 318–324 (2009).

37. Gulde, P., Hughes, C. M. L. & Hermsdörfer, J. Effects of Stroke on Ipsilesional End-Effector Kinematics in a Multi-Step Activity of Daily Living. Front. Hum. Neurosci. 11, (2017).

38. Dayan, P. & Balleine, B. W. Reward, motivation, and reinforcement learning. Neuron 36, 285–298 (2002).

39. Sutton, R. S. & Barto, A. G. Reinforcement Learning: An Introduction. 352.

40. Manohar, S. G. et al. Reward Pays the Cost of Noise Reduction in Motor and Cognitive Control. Current Biology 25, 1707 (2015).

41. Codol, O., Holland, P. J., Manohar, S. G. & Galea, J. M. Reward-based improvements in motor control are driven by multiple error-reducing mechanisms. J. Neurosci. (2020) doi: 10.1523/JNEUROSCI.2646-19.2020.

42. Summerside, E. M., Shadmehr, R. & Ahmed, A. A. Control of Movement: Vigor of reaching movements: reward discounts the cost of effort. Journal of Neurophysiology 119, 2347 (2018).

43. Codol, O., Galea, J. M., Jalali, R. & Holland, P. J. Reward-driven enhancements in motor control are robust to TMS manipulation. Exp Brain Res 1–13 (2020) doi: 10.1007/s00221-020-05802-1.

44. Kawagoe, R., Takikawa, Y. & Hikosaka, O. Reward-Predicting Activity of Dopamine and Caudate Neurons—A Possible Mechanism of Motivational Control of Saccadic Eye Movement. Journal of Neurophysiology 91, 1013–1024 (2004).

45. Pekny, S. E., Izawa, J. & Shadmehr, R. Reward-Dependent Modulation of Movement Variability. The Journal of Neuroscience 35, 4015 (2015).

46. Galea, J. M., Mallia, E., Rothwell, J. & Diedrichsen, J. The dissociable effects of punishment and reward on motor learning. Nat Neurosci 18, 597–602 (2015).

47. Carroll, T. J., McNamee, D., Ingram, J. N. & Wolpert, D. M. Rapid Visuomotor Responses Reflect Value-Based Decisions. The Journal of Neuroscience 39, 3906 (2019).

48. Goodman, R. N. et al. Increased reward in ankle robotics training enhances motor control and cortical efficiency in stroke. J Rehabil Res Dev 51, 213–227 (2014).

49. Quattrocchi, G., Greenwood, R., Rothwell, J. C., Galea, J. M. & Bestmann, S. Reward and punishment enhance motor adaptation in stroke. J. Neurol. Neurosurg. Psychiatry 88, 730–736 (2017).

50. Abe, M. et al. Reward improves long-term retention of a motor memory through induction of offline memory gains. Curr. Biol. 21, 557–562 (2011).

51. Wächter, T., Lungu, O. V., Liu, T., Willingham, D. T. & Ashe, J. Differential effect of reward and punishment on procedural learning. J. Neurosci. 29, 436–443 (2009).

52. Steel, A., Silson, E. H., Stagg, C. J. & Baker, C. I. The impact of reward and punishment on skill learning depends on task demands. Sci Rep 6, (2016).

53. Flash, T. & Hogan, N. The coordination of arm movements: an experimentally confirmed mathematical model. J. Neurosci. 5, 1688–1703 (1985).

54. Huang, H. J., Kram, R. & Ahmed, A. A. Reduction of Metabolic Cost during Motor Learning of Arm Reaching Dynamics. Journal of Neuroscience 32, 2182–2190 (2012).

55. Balasubramanian, S., Melendez-Calderon, A., Roby-Brami, A. & Burdet, E. On the analysis of movement smoothness. Journal of NeuroEngineering and Rehabilitation 12, 112 (2015).

56. Gribble, P. L., Mullin, L. I., Cothros, N. & Mattar, A. Role of cocontraction in arm movement accuracy. J. Neurophysiol. 89, 2396–2405 (2003).

57. Dhawale, A. K., Smith, M. A. & Ölveczky, B. P. The Role of Variability in Motor Learning. Annu Rev Neurosci 40, 479–498 (2017).

58. S, D. I’ (y) h m h m: v b y, x k. Current Opinion in Behavioral Sciences 20, 183–195 (2018).

59. Shmuelof, L., Krakauer, J. W. & Mazzoni, P. How is a motor skill learned? Change and invariance at the levels of task success and trajectory control. J Neurophysiol 108, 578–594 (2012).

60. Schmidt, R. A. & Lee, T. D. Motor control and learning: A behavioral emphasis, 5th ed. ix, 581 (Human Kinetics, 2011).

61. Kornysheva, K. et al. Neural Competitive Queuing of Ordinal Structure Underlies Skilled Sequential Action. Neuron 101, 1166-1180.e3 (2019).

62. Santos, F. J., Oliveira, R. F., Jin, X. & Costa, R. M. Corticostriatal dynamics encode the refinement of specific behavioral variability during skill learning. Elife 4, e09423 (2015).

63. Collins, A. L. et al. Dynamic mesolimbic dopamine signaling during action sequence learning and expectation violation. Sci Rep 6, 20231 (2016).

64. https://www.mathworks.com/matlabcentral/mlc-downloads/downloads/submissions/58403/versions/7/previews/min_jerk_todorov.m/index.html.

65. Gulde, P. & Hermsdörfer, J. Smoothness Metrics in Complex Movement Tasks. Front. Neurol. 9, (2018).

66. siva82kb/smoothness. GitHub https://github.com/siva82kb/smoothness.

67. Sadnicka, A. et al. High motor variability in DYT1 dystonia is associated with impaired visuomotor adaptation. Scientific Reports 8, 1–11 (2018).

68. Revol, P. et al. Pointing errors in immediate and delayed conditions in unilateral optic ataxia. Spat Vis 16, 347–364 (2003).

69. D’E c, J. SLM - Shape Language Modeling. https://de.mathworks.com/matlabcentral/fileexchange/24443-slm-shape-language-modeling (2020).

